# Expectation violations produce error signals in mouse V1

**DOI:** 10.1101/2021.12.31.474652

**Authors:** Byron H. Price, Cambria M. Jensen, Anthony A. Khoudary, Jeffrey P. Gavornik

## Abstract

Repeated exposure to visual sequences changes the form of evoked activity in the primary visual cortex (V1). Predictive coding theory provides a potential explanation for this, namely that plasticity shapes cortical circuits to encode spatiotemporal predictions and that subsequent responses are modulated by the degree to which actual inputs match these expectations. Here we use a recently developed statistical modeling technique called Model-Based Targeted Dimensionality Reduction (MbTDR) to study visually-evoked dynamics in mouse V1 in context of a previously described experimental paradigm called “sequence learning”. We report that evoked spiking activity changed significantly with training, in a manner generally consistent with the predictive coding framework. Neural responses to expected stimuli were suppressed in a late window (100-150ms) after stimulus onset following training, while responses to novel stimuli were not. Omitting predictable stimuli led to increased firing at the expected time of stimulus onset, but only in trained mice. Substituting a novel stimulus for a familiar one led to changes in firing that persisted for at least 300ms. In addition, we show that spiking data can be used to accurately decode time within the sequence. Our findings are consistent with the idea that plasticity in early visual circuits is involved in coding spatiotemporal information.

## Introduction

Sensory inputs contain statistical regularities that convey information about the external environment. Numerous lines of experimental and theoretical evidence suggest that the brain exploits these regularities to understand and predict the structure of the world around us (H. B. Barlow, 1961; Chalk, Marre, & Tkačik, 2018; Friston, 2005; Hosoya, Baccus, & Meister, 2005; Keller & Mrsic-Flogel, 2018; Palmer, Marre, Berry, & Bialek, 2015; Rao & Ballard, 1999; Spratling, 2017; Zmarz & Keller, 2016). Such predictions are useful for a wide variety of behavioral and neural tasks including efficient movement coordination (Körding & Wolpert, 2004; McNamee & Wolpert, 2019; Wolpert, Ghahramani, & Jordan, 1995), trajectory extrapolation (Montague & Sejnowski, 1994; Palmer et al., 2015), and information compression (H. Barlow, 2001a, 2001b; Dan, Atick, & Reid, 1996; Srinivasan, Laughlin, & Dubs, 1982) among others. Exactly how this is done remains an open question, but it is becoming increasingly clear that early visual areas encode efficient representations of their inputs both in space and across time (H. Barlow, 2001b; H. B. Barlow, 1961; Collewijn et al., 2008; Dan et al., 1996; Elias, 1955; Hosoya et al., 2005; Kelly, 1985; Kuang, Poletti, Victor, & Rucci, 2012; Lappe, Bremmer, & Van Den Berg, 1999; Leinweber, Ward, Sobczak, Attinger, & Keller, 2017; Palmer et al., 2015; Rucci, 2008; Srinivasan et al., 1982; Zmarz & Keller, 2016). The predictive coding model posits that this is accomplished by comparing predictions against incoming sensory data, then transmitting prediction error signals (Elias, 1955; Keller & Mrsic-Flogel, 2018; Rao & Sejnowski, 2001; Spratling, 2017; Srinivasan et al., 1982; Zmarz & Keller, 2016). This model has many attractive elements, and may help explain phenomena ranging from extra-classical receptive field properties in V1 (Atick & Redlich, 1993; Rao & Ballard, 1999) to the activity of dopamine neurons in the ventral tegmental area (VTA) (W. Schultz, Dayan, & Montague, 1997; Wolfram Schultz, 2016). However, many biological implementation details remain vague and specific elements of the model have not been validated in the brain. Particularly lacking is a clear explanation of how experience-dependent plasticity encodes the temporal relationships required for predictions into neural circuits.

Experimental evidence originating in multiple labs over the last decade has demonstrated that visual experience shapes V1 circuits to encode and predict temporal relationships (Finnie, Komorowski, & Bear, 2021; Garrett et al., 2020; Gavornik & Bear, 2014; Gillon et al., 2021; Homann, Koay, Glidden, Tank, & Berry, 2017; Orbán, Berkes, Fiser, & Lengyel, 2016; Shuler & Bear, 2006; Weliky, Fiser, Hunt, & Wagner, 2003; Zmarz & Keller, 2016). Insights gained from V1 may apply broadly to other regions as well (Douglas & Martin, 2004; Douglas, Martin, & Whitteridge, 1989; Edelman & Mountcastle, 1978; Hawkins & Ahmad, 2016), making this an experimentally accessible area in which to study the cortical basis of predictive processing. In the present study, we investigated predictive processing in the context of a specific form of sequence learning previously described as in V1 (Finnie et al., 2021; Gavornik & Bear, 2014) and anterior cingulate cortex (ACC) (Sidorov et al., 2020). Specifically, we passively exposed mice to rapidly-flashed sequences of sinusoidal gratings, whose spatiotemporal structure differs dramatically from the approximate 1/*f* spectrum of their natural environment (Carandini et al., 2005; Ocko, Lindsey, Ganguli, & Deny, 2018; Olshausen & Field, 1997). In doing so, we sought to determine whether prediction errors were generated in V1 and the extent to which those prediction errors were tuned to the spatiotemporal structure of the learned sequence.

## Results

### Experimental Design & Sequence Stimulus

Our goal in designing the experiment was to characterize how neural responses change with experience, under the assumption that we could not track individual neurons across days. To do so we created a novel randomized Train-Test experimental protocol (Figure 1) that presented a set of Test stimuli to mice after zero to three days of training. During training, each mouse was presented with a with a single training sequence, ABCD (Figure 1d), in order to condition them to expect the specifical spatiotemporal structure of that sequence. After training, mice viewed randomized test sequences to determine how different types of expectation violation change neural responses in V1 (Figure 1e). Animals were randomly assigned a Test day (1-4). Animals that saw the Test set on Day 1 (e.g. after zero days of training) served as naïve controls for the “trained” mice who saw the Test stimulus after at least one day of training. Each mouse saw the Test stimulus set only once and then were removed from the experiment. The randomization procedure insured that the amount of data collected from each Test day group was approximately balanced (Figure 1b-c).

**Figure 1:**
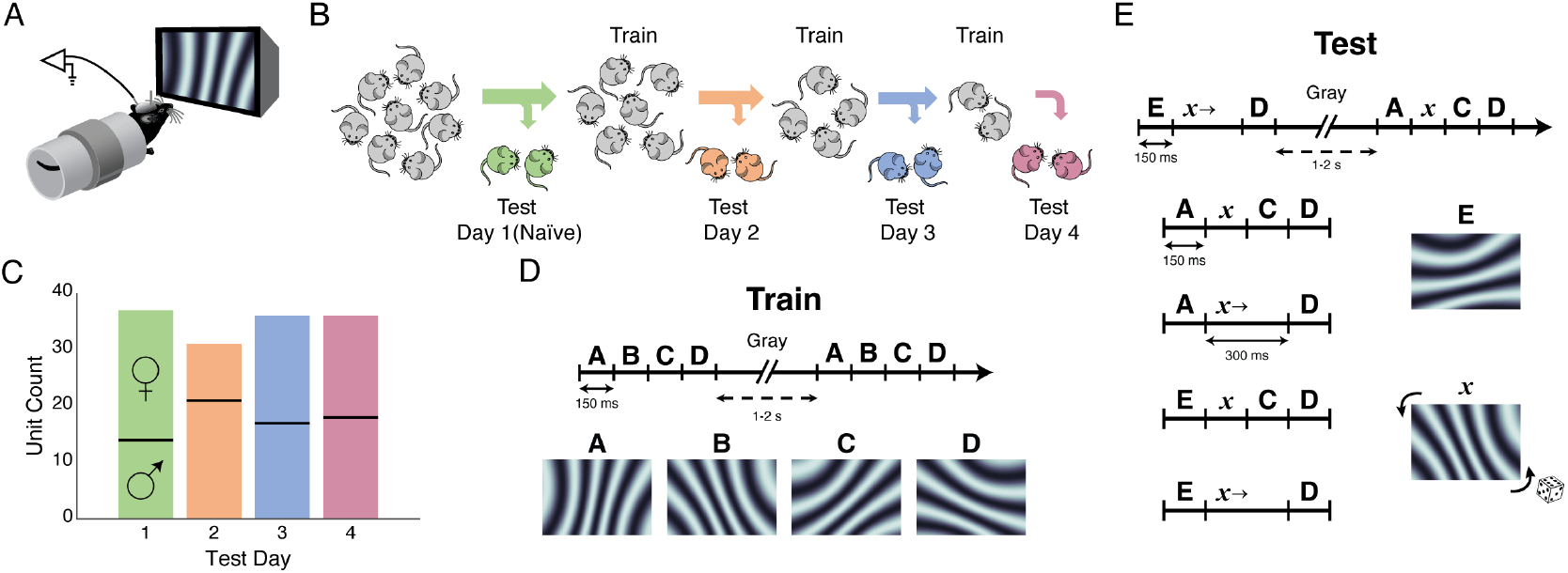
Experimental Design. **A**. Awake head-fixed mice viewed stimuli from a distance of 25 cm simultaneous with extracellular recoding via chronically implanted wire bundles. **B**. Mice were randomly assigned to see Test stimuli on day 1 (25%, naïve controls, green), day 2 (33% of remaining mice, orange, 1 day of training), day 3 (50% of remaining, blue, 2 days of training), or day 4 (100% of remaining, magenta, 3 days of training). **C**. The number of units recorded on each day was approximately uniform and evenly distributed between female and male mice (top & bottom of each bar). In total, we recorded 140 unique multi-unit channels from 56 mice. **D**. Each training session consisted of 200 presentations of the sequence ABCD. Each element had a unique orientation and was held on screen for 150ms and the full sequence lasted 600 ms. Sequences were separated from each other by a gray screen, held for a uniform random interval between 1 and 2 seconds. **E**. During Test sessions mice were exposed to 600 presentations of randomly selected novel sequences: AxCD, ExCD, Ax→D, and Ex→D where x indicates a random orientation (uniformly distributed ± 60° around B) and → indicates an omitted third element (second element on screen for 300 ms, 50% of trials).

All mice were awake and head fixed while viewing sequence stimuli during both *Train* and *Test* sessions. Each element in the sequence lasted 150 ms and a complete 4-element sequence lasted 600 ms (Figure 1d,e). Element transitions within a sequence were continuous and a gray screen was used to separate individual sequence presentations. To eliminate the possibility that the animal could anticipate when the next sequence would begin, the gray screen was displayed for a random interval drawn from a uniform distribution between 1 and 2 seconds. In all sessions (*Train* and *Test*), we recorded spiking data from binocular V1 Layer 4 neurons (Supplemental Figure 1), along with 50-Hz infrared video of the face and whisker pad (Supplemental Figure 2). In total, we identified 140 distinct, visually-responsive multi-unit channels from 56 mice (29 females, 27 males; P66.8 +/- 0.5 days at experimental start [mean +/- SEM]; see Methods for unit inclusion criteria).

During *Test* sessions, each sequence was initiated with the familiar element A or a novel element E that was not contained in the training sequence. The second *Test* sequence element, *x*, was a randomly oriented grating held on the screen for either 150 ms (A*x*CD and E*x*CD) or, in 50% of trials, 300 ms (A*x*→D and E*x*→D). This design allowed us to test the importance of familiarity (A vs E), orientation (*x*), temporal expectation (→), and training as well as to look for evidence of error signals associated with positive (unexpected inclusion) and negative (unexpected omission) prediction errors as described in various predictive coding models (Keller & Mrsic-Flogel, 2018; Spratling, 2017).

Previous work has shown that stimulus-evoked movements can change with V1 plasticity (Cooke & Bear, 2015), so we analyzed the video data to determine whether a similar effect occurs following sequence learning. Though neural variability correlated with movement generally, as has been reported elsewhere, we found no evidence for a relationship between the mouse’s movement and the timing of visual stimulation (Supplemental Figure 2). Therefore, we ignored the movement data in all subsequent analyses.

### A Statistical Model (MbTDR) Captures Stimulus-Dependent Neural Variability

We used a statistical modeling tool known as model-based targeted dimensionality reduction (MbTDR) (Aoi, Mante, & Pillow, 2020; Aoi & Pillow, 2018) to analyze our high dimensional, heterogeneous data. MbTDR is a supervised probabilistic model that projects high-dimensional data onto low-rank subspaces spanned by different regression covariates (Figure 2a). Each subspace is a set of shared basis functions, low-dimensional neural trajectories, along with “unit factors” that specify how much each basis contributes to the trial-by-trial firing rate of a given unit (Figure 2a). We created covariates related to the structure of the sequence stimulus and our experimental design, including condition independent (baseline PSTHs), experimental day, first element (A or E, encoded as an indicator function, [*I*(*E*)]), the orientation of the randomized second element (*Angle*(*x*) & *Angle*(*x*)^2^), second element duration [*I*(*x*→)], trial number, and combinations/interaction-terms thereof. Using a greedy forward stepwise algorithm that minimizes the Akaike Information Criterion, the model selects the optimal rank for each covariate and interaction term. The algorithm allows the optimal rank for a given covariate to be zero and finding a rank greater than zero is comparable to discovering a significant effect for that covariate in a regression model such as ANOVA. The optimal rank is the number of unique PSTH templates that are necessary to explain the variability in PSTHs across the population. If the responses for all units were completely independent, and no low-dimensional representation were possible, then the optimal rank for each covariate would equal the total number of recorded units.

**Figure 2:**
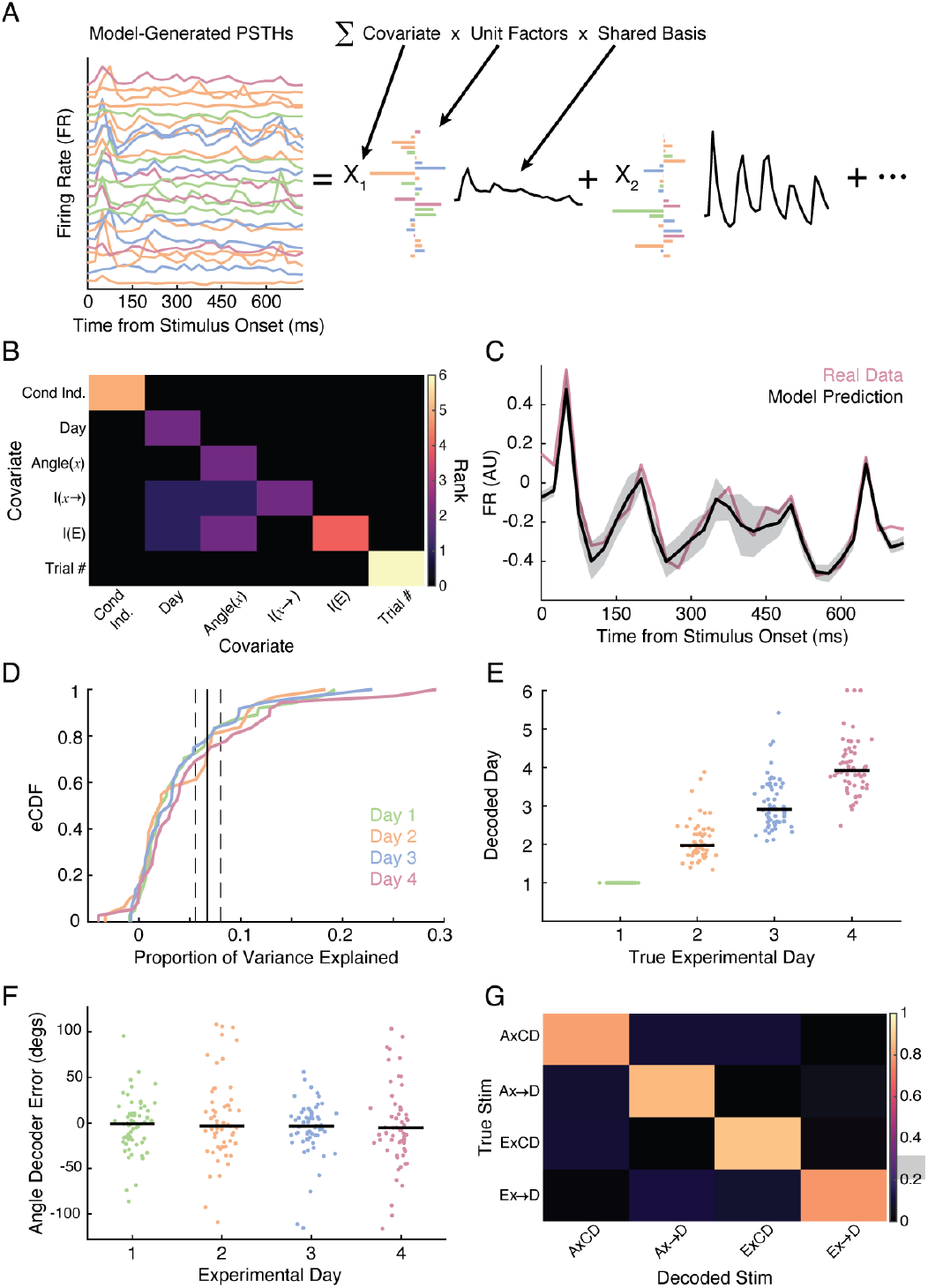
MbTDR Captures Stimulus-Dependent Neural Variability. **A**. Model-based targeted dimensionality reduction (MbTDR, after Aoi & Pillow 2018) models PSTHs as a linear combination of covariates, unit factors, and shared basis functions. MbTDR discovers a low-rank representation, consisting of a small set of shared bases that are weighted differently for each unit (color code as in Figure 1b). **B**. Rank of optimal model, fit by maximum likelihood and a greedy rank estimation algorithm. Covariates were chosen to represent the experimental design and Test stimulus (see main text for details). Diagonal elements are the rank of each covariate, and off-diagonal elements represent the rank of interaction terms. **C**. An example day-4-unit PSTH (binned at 25ms with no smoothing, z-scored to zero spontaneous baseline firing and unit variance, as in all subsequent figures and analyses). The MbTDR prediction (black) matches the neural data (magenta, 95% confidence interval computed from observed Fisher information in gray). This unit had 4.41% held-out explained variance. **D**. Empirical cumulative distribution functions (eCDFs) of explained variance across the 140 units. Each line represents the eCDF computed using held-out data from one Test day (60 trials / day). The vertical black line is an estimate of explainable variance with a bootstrap 95% confidence interval. **E**. Test days can be accurately decoded (each dot represents a single held-out pseudo-trial, x-axis jitter added for visibility). Due to our experiment and model design, Day 1 trials are automatically known. **F**. Decoding error for x, the randomized angle of the second element, on the same held-out data (each dot is one pseudo-trial). The standard deviation on the error is 38.7 degrees. **G**. Confusion matrix for decoding the four primary stimulus types. Overall accuracy was 82.5%, while chance for 60 trials is 25 +/- 2.7% (mean +/- SEM, gray box on color bar shows 95% confidence interval of this estimate). The identity matrix would constitute perfect accuracy.

We fit a single model for all recorded units on 90% of trials using maximum likelihood and the greedy algorithm mentioned above; 10% of the data was held-out of the model fitting in order to estimate explained variance and to use for decoding. The final total rank of the model was 26, with ranks for each covariate and interaction term displayed in Figure 2b (model bases are depicted in Supplemental Figure 5). All of the fundamental covariates were significant, but the only significant interaction terms were: *I*(*x*→) * *Day, I*(*E*) * *Day, I*(*x*→) * *Angle*(*x*), *I*(*E*) * *Angle*(*x*), *I*(*E*) * *Angle*(*x*)^2^. The relevance of these terms is discussed below (for more details on the model and covariates, see Methods and Supplemental Table 1).

The final model achieved 6.2% explained variance on the held-out data, which was comparable to our estimate of 6.7 +/- 0.7% *explainable variance* (mean +/- SEM) (Figure 2c-d). In order to estimate explainable variance, we fit PSTHs to repeated presentations of the same sequence (ABCD on Training days). In this case, the PSTH captures predictable variability due to the stimulus, and all unexplained variance must be due to factors that are inconsistent across trials (movement, neural variability, etc.). There was a strong positive correlation between evoked firing rate and held-out explained variance, despite the fact that units were normalized to unit variance prior to model fitting: *ρ* = 0.45 (p<1e-6, Spearman’s rho permutation test, N=140 units). However, there was no such correlation between explained variance and experimental day: *ρ* = 0.053 (p=0.54), indicating that model fit was comparable regardless of the number of training days. Finally, 125/140 units had greater than 0% held-out explained variance, and 48/140 had greater than 5% (see Figure 2c for an example PSTH from a unit with about 5% explained variance), indicating the model accurately captured stimulus-dependent, trial-by-trial fluctuations in neural firing.

To further convince ourselves the model fit the data well, we used Bayes’ theorem to decode stimulus covariates on the held-out data (Figure 2e-g). For each *Test* day, we created 60 pseudo-trials by combining data from all recorded units, about 35 per day. Stimulus features could be accurately decoded on individual pseudo-trials, revealing strong correlations with ground truth: *ρ* = 0.78 for experimental day (p<1e-6, Spearman’s rho permutation test, excluding Day 1, Figure 2e); *ρ* = 0.50 for angle of *x* (p<1e-6, Figure 2f); *ρ* = 0.59 for trial number (p<1e-6); *ρ* = 0.80 for the third element omitted (*I*(*x*→), p<1e-6); and *ρ* = 0.81 for E starting the sequence (*I*(*E*), p<1e-6). When asking the decoder to determine which of the four primary Test stimulus types had been displayed (A*x*CD, E*x*CD, A*x*→D, or E*x*→D), the decoder achieved 82.5% accuracy on 240 held-out pseudo-trials, compared to 25 +/- 2.8% by chance (Figure 2g). Together, these results demonstrate the efficacy of the model in capturing stimulus-dependent neural variability.

### MbTDR Reveals Coordinated Training-Dependent Changes in Neural Activity

We found a modest increase in evoked firing rate across Test days. The global median evoked firing rate was 7.8 Hz, while for each Test day it was: 1-6.4 Hz; 2-11.0 Hz; 3-10.6 Hz; 4-14.2 Hz (two-sided permutation test for difference in median firing rate: p = 0.026 Day 1 vs. Day 2; p = 0.034 for Day 1 vs. Day 3; p = 0.006 for Day 1 vs. Day 4). Besides this simple measure of learning-dependent change, the MbTDR model-selection process also chose *Day* as a significant covariate with a rank of 2. Units that were strongly modulated by these bases are shown in Figure 3a. Note that in this and subsequent figures firing rates are z-scored to zero spontaneous baseline firing and unit variance.

**Figure 3:**
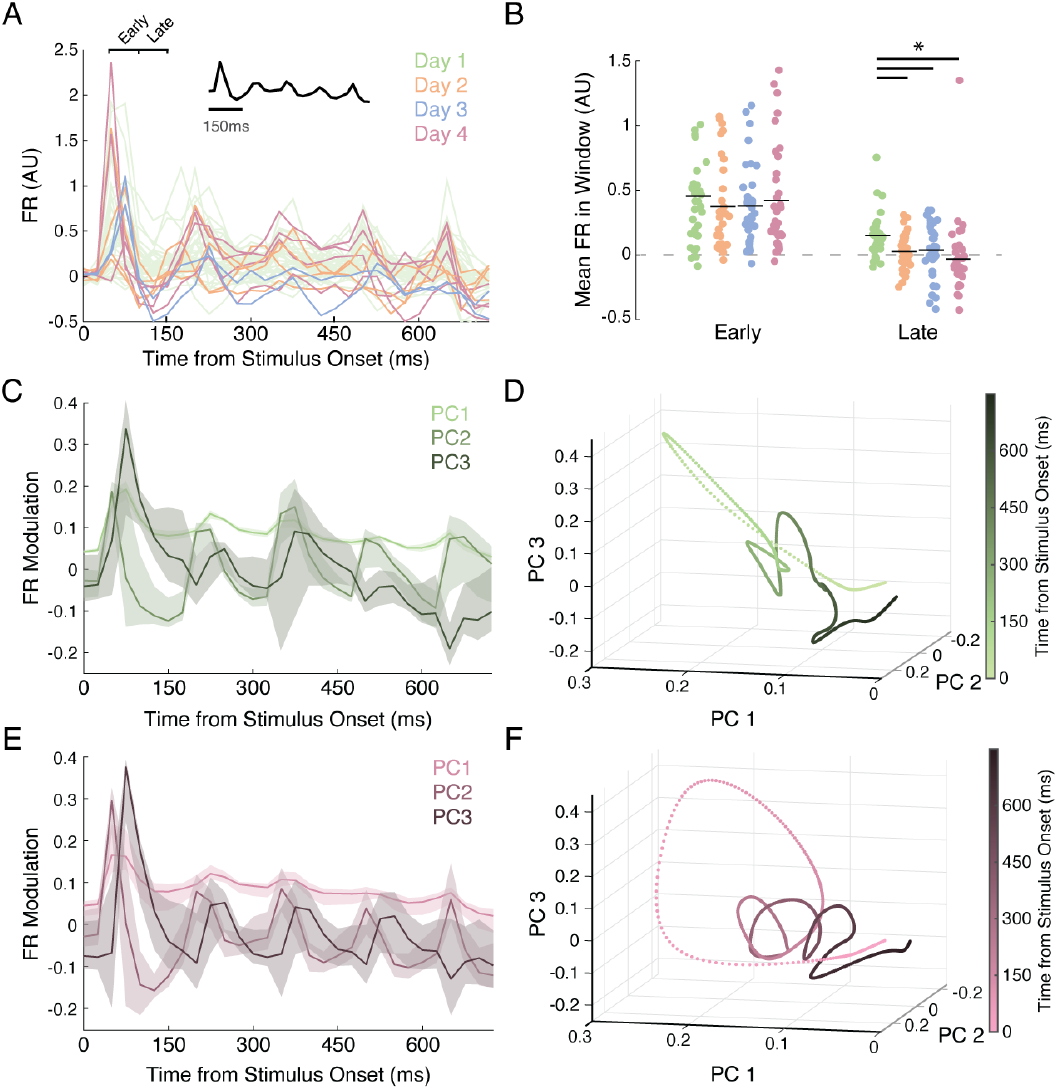
MbTDR Reveals Coordinated Training-Dependent Change in Neural Activity. **A**. Example PSTHs to AxCD (computed from the raw data, not the model fit). Units from trained mice (Days 2,3,4) that were strongly modulated by the Day covariate (inset) are overlaid above example PSTHs from Day 1 units (green). Note the dip in firing after the onset of A in trained units. Some trained units drop as low as -0.5 on this z-scale, while no unit from Day 1 (the naïve group) drops below -0.15 in that same window. Early / Late designations indicate the time windows used in B. **B**. Comparison of mean normalized firing rate across Test days (again computed from the raw data, not model fit). Left shows mean firing rate in an early window (51-100ms after onset of A), right during a late window (101-150ms). There was no significant difference between trained and naïve groups in the early window (two-sided permutation test for difference in mean from Day 1, with a threshold set to p<0.05/6=0.0083 for multiple comparisons: Day 2 p=0.441; Day 3 p=0.437; Day 4 p=0.668). However, evoked firing in the late window was significantly lower in all trained groups (**Day 2 p=6.92e-4; Day 3 p=0.0052; Day 4 p=5.02e-4**). Black dotted line marks the spontaneous baseline firing rate. **C**. First, second, and third principal components of evoked neural activity to ABCD in naïve mice, computed from the MbTDR fit (see Methods) with 95% confidence intervals. **D**. Dynamic latent trajectory representation of the data in C in principle component space. Data was projected from 25-ms bins as in C to 1-ms bins using a Gaussian radial basis with 12.5-ms standard deviation. **E**. Same as C, but for the set of Day 4 units. The first component remains unchanged across days, while the second and third differ dramatically from Day 1. Note the dip in the second component from Day 4 around 100ms, which mirrors decreased firing in the late window observed in B. **F**. Same as D, but for Day 4 units. The second and third dimensions show rotational dynamics which, along with the first component, create a spiraling latent trajectory.

Comparing evoked responses across days, it was visually evident that the overall increase in firing rate was not uniform over the duration of the sequence. The initial response following element transitions tended to increase in magnitude, while firing rate during a later sustained response decreased. To quantify this effect we calculated the average firing rates in early (51-100ms) and late (101-150ms) windows after the onset of element A across test days. While increases in firing during the early window were not significant, there was significantly less firing in the late window for units from trained mice (Figure 3b). This dip in firing was also captured by the MbTDR *Day* covariate, as seen in Figure 3a.

Next we used MbTDR to visualize low-dimensional dynamic neural trajectories across Test days by performing a singular-value decomposition of the model fits for all possible test sequences (see Methods). The first principal component (Figure 3c,e in light shade) shows an initial excursion at the onset of the stimulus and then a slow relaxation back to baseline. This was completely unchanged across days and accounted for about 91.7% of the variance in the PSTHs (i.e. signal variance) on Day 1 and 78.1% on Day 4. However, the second and third components, accounting for 6.2% of the variance on Day 1 and 16.5% on Day 4, changed dramatically between naïve and trained mice (Figure 3c-f). Figure 3c shows the first three components on Day 1, while Figure 3d provides a dynamic representation of those components plotted against one another. The dynamic representation from naïve mice crosses over itself and exhibits sharp direction changes, which might be described as highly tangled (Russo et al., 2018) or highly curved (Henaff et al., 2021; Hénaff, Goris, & Simoncelli, 2019). Fully trained mice, by comparison, evidenced stronger rotational dynamics in the second and third dimensions, and a spiraling trajectory in the first three components (Figure 3e-f).

### Orientation Tuning Does Not Shift Significantly with Training

We hypothesized that sequence learning would cause a shift in population orientation tuning from an approximately uniform distribution (all angles equally represented with some over-representation for the cardinal angles) toward a distribution with more mass on the orientations of the trained elements, ABCD. This kind of unsupervised shift in orientation tuning has been observed in other experimental protocols (Kaneko, Fu, & Stryker, 2017). Randomizing the angle of the second element allowed us to estimate orientation tuning curves for each unit and compare against the trained B. The MbTDR model-selection process picked out *Angle*(*x*), *Angle*(*x*)^2^, *I*(*x*→) * *Angle*(*x*), *I*(*E*) * *Angle*(*x*), and *I*(*E*) * *Angle*(*x*)^2^ as significant covariates. This suggests individual units were not only tuned to the orientation of the second element, but that their tuning depended on the identity of the first element (stimulus-history-dependent tuning). Supplemental Figure 3a shows the shared bases for the *Angle*(*x*) and *Angle*(*x*)^2^ covariates. From the MbTDR output, we can obtain a complete orientation tuning curve for each unit, along with a peak tuning angle (Suppl. Figure 3b). With these peak tuning values, we then constructed an empirical cumulative distribution function (populating tuning curve) for each Test day (Suppl. Figure 3c). The peak-tuning cumulative distribution functions were *not* significantly different between naïve and trained mice (two-sided KS test, naïve [Day 1] vs. trained [Days 3 & 4]: D=0.106, p=0.932). This result suggests that training does not shift second-element orientation tuning curves towards the trained orientation of B on a large scale, though we cannot rule out the possibility that some neurons shift. This question could be addressed with a longitudinal dataset tracking individual units across days.

### Unexpected Omissions Cause Negative Prediction Errors

Test-day stimuli A*x*→D and E*x*→D deviate from expectation by omitting the third element, C, from the sequence. This omission of an expected stimulus element was designed to elicit a negative prediction error in trained mice, indicating the absence of an expected stimulus. In units from naïve mice, neural firing during an omission reliably decayed back to baseline (red trace in Figure 4a for one example unit). In trained mice, by comparison, about 25% of units showed a response consistent with a negative prediction error (red trace in Figure 4b for example unit). In those units, firing rates ramped in anticipation of the expected onset of element C, reaching their peak at about 300-325ms after the onset of the sequence. When C was shown, its onset precipitated a rapid decrease in firing (blue trace in Figure 4b). When C was omitted, however, elevated firing persisted until the onset of the next sequence element, D (red trace in Figure 4b), consistent with a negative prediction error.

**Figure 4:**
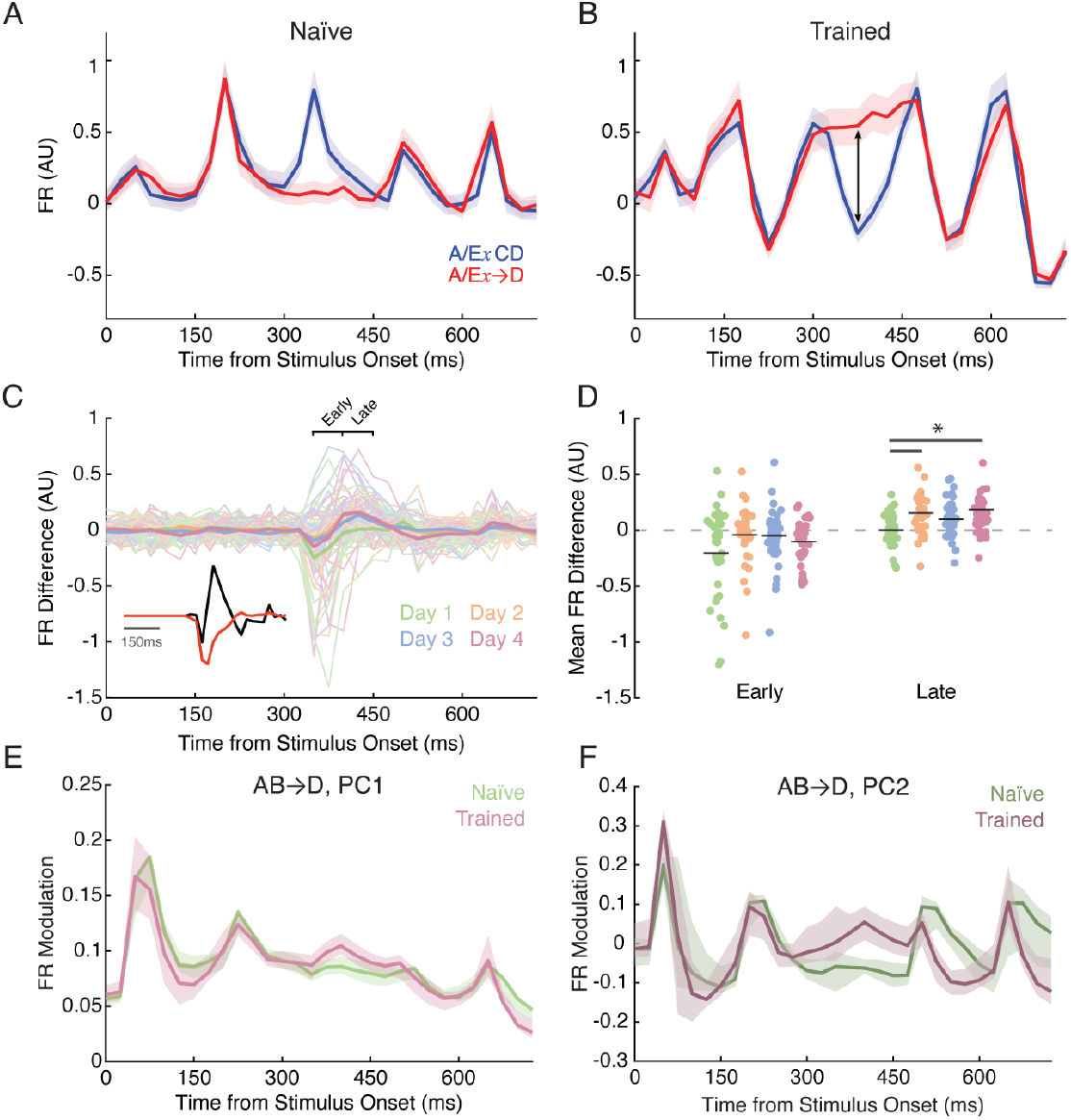
Unexpected Omissions Cause Negative Prediction Errors. **A**. Example PSTHs from a Day 1 / naïve unit comparing A/ExCD (blue) and A/Ex→D (red) trials (mean with 95% confidence intervals). **B**. Example PSTH from Day 3. Omission of element C (in red) drives increased and sustained firing in a late window after its expected onset. **C**. Difference PSTHs for all recorded units (red minus blue from previous example, marked by arrow in B). Inset shows shared basis functions from MbTDR fit for the I(x→) covariate (red) and interaction term I(x→)*Day (black). **D**. Comparison of difference PSTHs in early (left) and late (right) windows (marked in C) after expected onset of element C. The early window is 51-100ms after expected time of C onset, while late is 101-150ms. There was no significant difference between trained and naïve groups in the early window (two-sided permutation test for difference in mean from Day 1, with a threshold set to p<0.05/6=0.0083 for multiple comparisons: Day 2 p=0.061; Day 3 p=0.091; Day 4 p=0.194). In the late window, however, firing was significantly higher on Days 2 and 4 (**Day 2 p=0.0012**; Day 3 p=0.016; **Day 4 p=5.02e-4**). **E-F**. The first and second principal components of neural activity for the AB→D condition, in naïve (green) and fully trained (magenta) mice with 95% confidence intervals.

To further investigate this phenomenon, we created a set of difference PSTHs for all recorded units (Figure 4c) by subtracting the blue traces in Figure 4a-b from the red for all units (A/E*x*→D – A/E*x*CD). The difference PSTHs clearly show a tendency for units from trained mice to fire at an elevated rate when C is omitted (Figure 4c, thick lines depict the mean across units, positive values indicate firing is greater when C is omitted). To quantify this effect, we calculated the mean firing rate difference in early (51-100ms) and late (101-150ms) windows after the expected onset of C (Figure 4d). While there was no significant difference in the early window, trained units from Days 2 and 4 had significantly higher firing-rate differences in the late window than those differences computed from naïve units. Defining all those units as prediction-error units whose late-window mean firing rate difference exceeded three standard deviations of the baseline, we found the following proportions on each day: Day 1-3/37 (8.1%); Day 2-7/31 (22.6%); Day 3-7/36 (19.4%), Day 4-13/36 (36.1%).

Furthermore, the MbTDR model-selection process chose *I*(*x*→) and *I*(*x*→) * *Day* as significant covariates (Figure 4c inset, also Figure 2b). The shared basis functions recapitulate the difference PSTHs, showing that only in trained units (black trace, Figure 4c inset) the omission of C led to increased firing in the late window. We also used the model fit to visualize the first and second principal components of neural activity (Figure 4e-f). Comparing Day 1 and Day 4, these principal components clearly provide evidence for a negative prediction error. On Day 4, for example, PC2 shows a sustained increase in firing after the expected onset of C, while on Day 1 PC2 recapitulates the example unit from Figure 4a by decaying to baseline.

We next sought to determine the spatiotemporal specificity of the negative prediction errors, i.e. the extent to which they are tuned to the precise sequence observed during training (ABCD). We hypothesized that angles similar to B would elicit larger negative prediction error responses than angles far from B, but only in trained mice. The model selection procedure chose *I*(*x*→) * *Angle*(*x*) as a significant interaction covariate, though it did not choose other covariates that might govern the orientation tuning of the negative prediction error response, such as *I*(*x*→) * *I*(*E*) or *I*(*x*→) * *Angle*(*x*) * *Day*. Therefore, the negative-prediction-error effect likely depends on the angle of the preceding element, but not the angle of the first element (A versus E). However, because of the design of the model, the *I*(*x*→) * *Angle*(*x*) term can have a complex relationship with other covariates. For example, the significant *Angle*(*x*)^2^ covariate implies that neurons are tuned quadratically to the orientation of the second element, as expected in V1. However, the underlying shared basis function for *Angle*(*x*)^2^ extends in time to at least 500ms into the sequence (see Supplemental Figures 3a, 4a). This implies that orientation tuning curves for the second element, *x*, may be different from negative-prediction-error tuning curves computed during the omitted third element. To investigate this further, we used the model fit to determine the orientation tuning of the negative prediction errors during the early and late windows (51-100ms and 101-150ms after the expected onset of C).

Supplemental Figure 4 depicts several example prediction error orientation tuning curves, from which we extracted peak tuning angles for each unit, as before. Across the population of recorded units, there was no appreciable difference between peak tuning angles in naïve and trained mice. We therefore conclude that there is limited spatiotemporal specificity to the negative prediction errors, i.e. they seem to occur even when the preceding visual sequence elements do not match those presented during training. This may reflect a limitation in the ability of layer 4 cells in V1 to adapt to the novel spatiotemporal statistics of the trained sequence, or perhaps that the step-function-like transitions between sequence elements are sufficiently salient to elicit the predictive response themselves.

### Unexpected Substitutions Cause Positive Prediction Errors

The *Test* day stimulus set was designed to induce not only negative prediction errors, but also positive prediction errors that occur when an expected stimulus is replaced by an unexpected one. Given that the mice always saw ABCD during Training, replacing the expected A with a novel E violates expectation and should produce positive prediction errors. The MbTDR model selection process chose *I*(*E*) and *I*(*E*) * *Day* as significant covariates. The *I*(*E*) covariate suggests that the neural responses to A and E are different, as expected due to their differing orientations. The *I*(*E*) * *Day* covariate suggests that this difference between A and E evolves with training (see Figure 5c inset for shared basis functions corresponding to these covariates).

**Figure 5:**
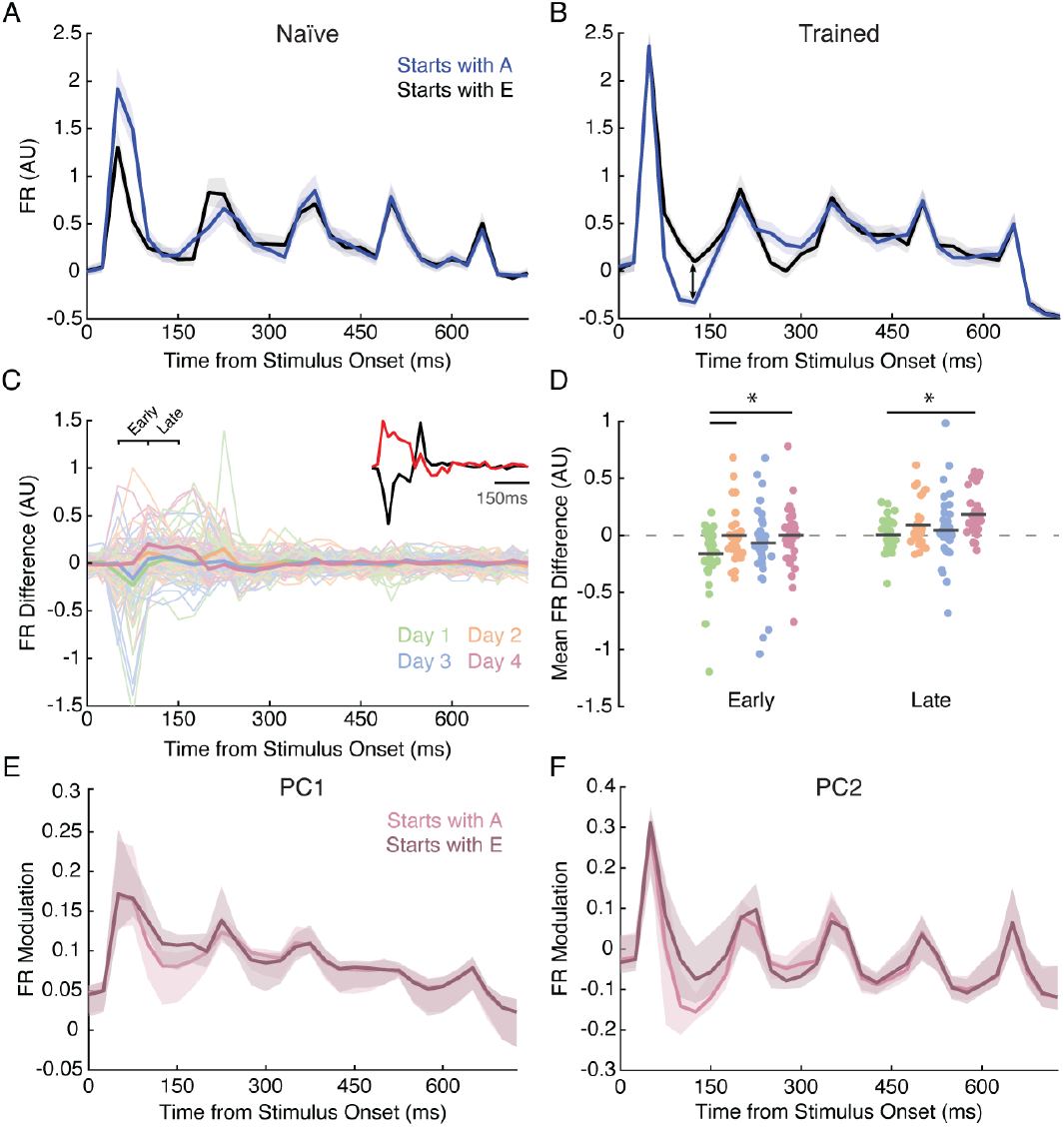
Unexpected Substitutions Cause Positive Prediction Errors. **A**. Example PSTHs from a Day 1 / naïve unit comparing all trials starting with A (blue) and E (black) with 95% confidence intervals. **B**. As in A but for a Day 4 unit. Presentation of the trained A drives a sustained decrease in firing during a late window after its onset that is not present following E. **C**. Difference PSTHs from all recorded units (black minus blue, illustrated with arrow in B). Inset shows shared basis functions from MbTDR fit for the I(E) covariate (black) and interaction term I(E)*Day (red). **D**. Comparison of difference PSTHs in early (left, 51-100ms) and late (right, 101-150ms) windows after onset of the first sequence element (A or E). There was a significant difference between naïve and trained groups in the early window (two-sided permutation test for difference in mean from Day 1, with a threshold set to p<0.05/6=0.0083 for multiple comparisons: **Day 2 p=0.0062**; Day 3 p=0.23; **Day 4 p=0.0071**). In the late window, the firing rate difference was significantly greater only on Day 4 (Day 2 p=0.029; Day 3 p=0.34; **Day 4 p=4.0e-6**). **E-F**. The first and second principal component of neural activity for the ABCD (light shade) and EBCD (dark shade) conditions with 95% confidence intervals. Both the first and second components show a significant dip following A, but not E, in the same late window.

In the PSTHs from trained mice, sequences starting with the familiar A showed decreased firing in a late window after stimulus onset (Figure 5b, also see Figure 3a,b). By comparison, sequences starting with E looked much more like the responses in naïve units (Figure 5b black trace, compared to Figure 5a black or blue). To quantify this effect, we looked at the firing rate differences between sequences starting with E and those starting with A (Figure 5c); these are the black traces from Figure 5a-b minus the blue. The firing rate differences in trained mice were significantly different from Day 1 in both the early and late windows (Figure 5d). During the late window in naïve mice, responses to A and E were approximately equal. In fully trained mice, however, the responses to A in the late window were significantly lower than those to E. This effect, of decreased firing in the late window for A but not E, was also picked up by MbTDR (Figure 5e,f). Training therefore created a differential neural response between familiar and novel stimuli. More precisely, the data suggests that training initiates a late-evolving inhibitory process that is only activated by expected stimuli. Unexpected stimuli, whether in naïve mice seeing the stimulus for the first time or in trained mice seeing a novel stimulus, do not activate the same process and show elevated firing in the late window.

During training, element A reliably predicted B. Assuming that learning encodes expected sequence order, the response to element B should look different if it is preceded by A (expected) versus E (unexpected). The model basis for *I*(*E*) modulates neural responses for about 400ms (supplemental Figure 5) supporting the idea that individual elements influence the responses to subsequent visual stimuli. To investigate further, we identified sequences on Test days where the angle of the second element *x* was within +/- 5 degrees of the trained element B, dubbed 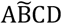 and 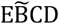. As shown in Figure 6a-b, the response to 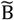 was different between naïve and trained mice only for the sequence 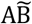. In the late window after the onset of 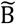, units from trained mice showed reduced firing relative to the naïve group, comparable to the positive prediction error effect observed between A and E. There was no significant difference between naïve and trained mice when E initiated the sequence. Though the statistics suggest that the response to the second element in the sequence depends on training and is differentiated by whether or not it is predicted by the preceding element, the data is not conclusive on this point. For example, the late-window 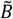 response following E shows a dip that looks similar to that seen when preceded by A. The current experimental design makes it difficult to interpret this result.

**Figure 6:**
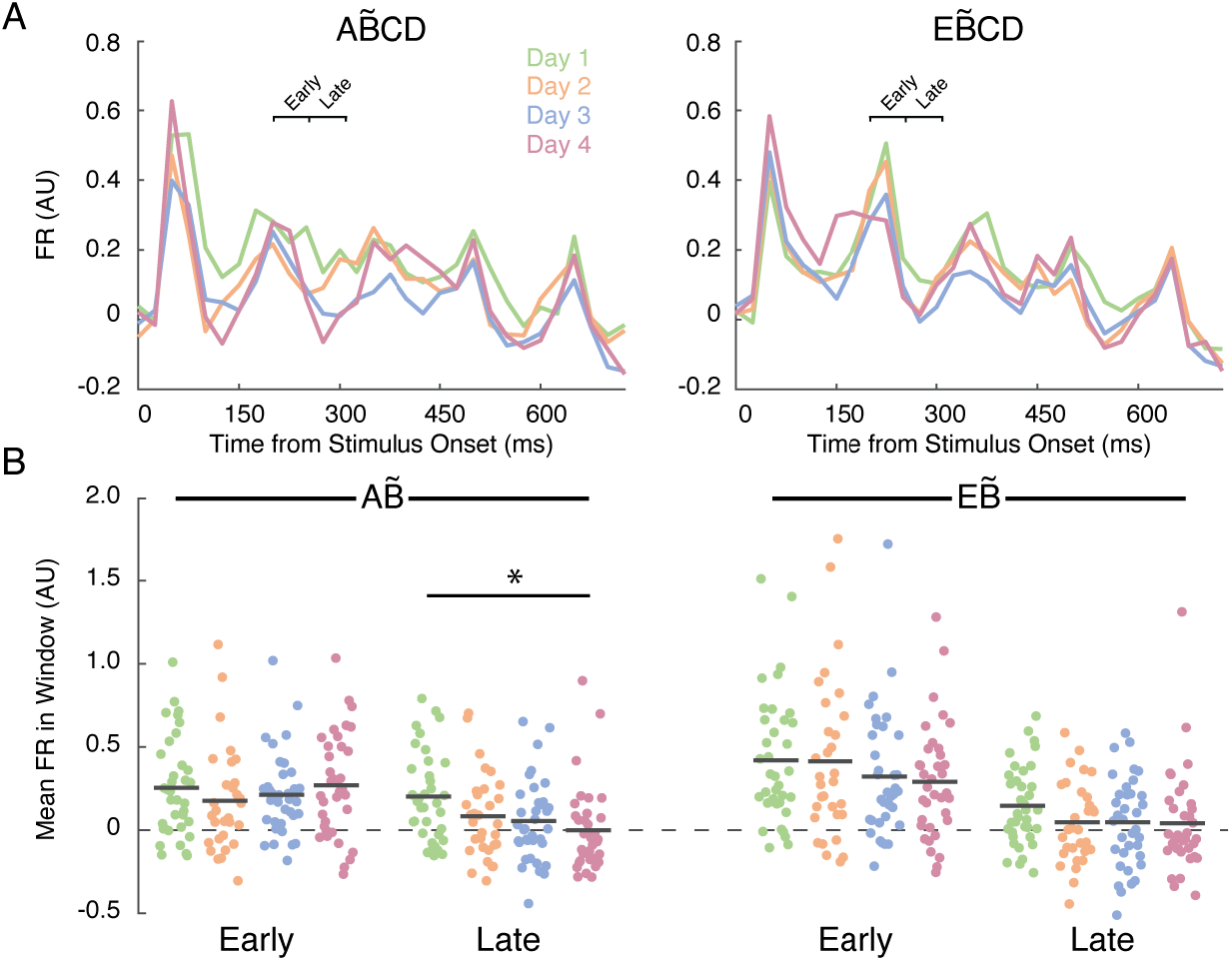
Predictions Span Element Transitions. **A**. Left: Average PSTHs across all recorded units to the sequence 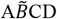, where 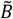 indicates that x was within +/- 5 degrees of the trained B (B was 120 degrees, so 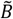 includes all angles from 115 to 125 degrees). Right: Same as Left, but for sequences 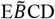 CD. **B**. Mean firing rate in early (51-100ms) and late (101-150ms, marked in A) windows following the onset of the second element, 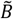, when preceded by A (left) or E (right). Each dot represents one unit. Only the late window for 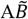 was significantly different between naïve and trained mice (two-sided permutation test for difference in mean from Day 1, with a threshold set to p<0.05/12=0.0042 for multiple comparisons: 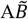 early Day 2 p=0.293; Day 3 p=0.520; Day 4 p=0.837. 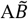 late Day 2 p=0.065; Day 3 p=0.016; **Day 4 p=0.001**. 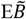 early Day 2 p=0.951; Day 3 p=0.280; Day 4 p=0.137. 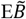 late Day 2 p=0.095; Day 3 p=0.100; Day 4 p=0.105).

### Reliable Temporal Information is Contained in the Neural Code

A prominent hypothesis in visual neuroscience proposes two distinct pathways for “what” and “where” information (Flindall & Gonzalez, 2020; Ungerleider & Mishkin, 1982). Goodale & Milner later proposed a “how” pathway, that does not simply locate objects in the environment but coordinates the sensorimotor transformations necessary to grab and manipulate them (Goodale & Milner, 1992; Milner & Goodale, 2008). In addition to these classical pathways, the early visual system may also be a prominent source of “when” information (Rauschecker, 2018). Such information could be useful to establish temporal context, predict the timing of future events, or coordinate sensorimotor behavior (Goel & Buonomano, 2014; Mauk & Buonomano, 2004). “When” information may be measured in relation to a saccade, the onset of a movement, or relative to co-occurring events. In the case of the sequence stimulus, the onset of the sequence marks a strong departure from ordinary visual experience and could therefore be used to initialize a clock. Based on these ideas, we hypothesized that training would improve our ability to decode elapsed time from the evoked neural population response.

To test our hypothesis, we used the MbTDR fit to perform instantaneous time decoding, attempting to use the model and held-out neural data to predict elapsed time on individual pseudo-trials and time bins (Figure 7a). The decoder takes in a neural population response vector from a single time bin and uses the model to predict which bin, of 30 possible, the data came from. If the neural code in V1 were a poor source of temporal information the decoder would fail to produce accurate time estimates. We expected the decoder to perform better with training and that is what we discovered (Figure 7c-d). On Day 1, the decoder achieved a soft accuracy of 26.0%, compared to 9.8 +/-1% by chance (soft accuracy allows for the decoder to be off by one time bin in either direction). On Day 2, soft accuracy was 24.9%; on Day 3, 29.8%; and on Day 4, 32.6%. We performed a two-sided permutation test on the difference in accuracy between Day 1 and all subsequent days, finding the accuracy to be significantly greater on Days 3 and 4 (Figure 7c). Thus, the ability to decode time improves with training.

**Figure 7:**
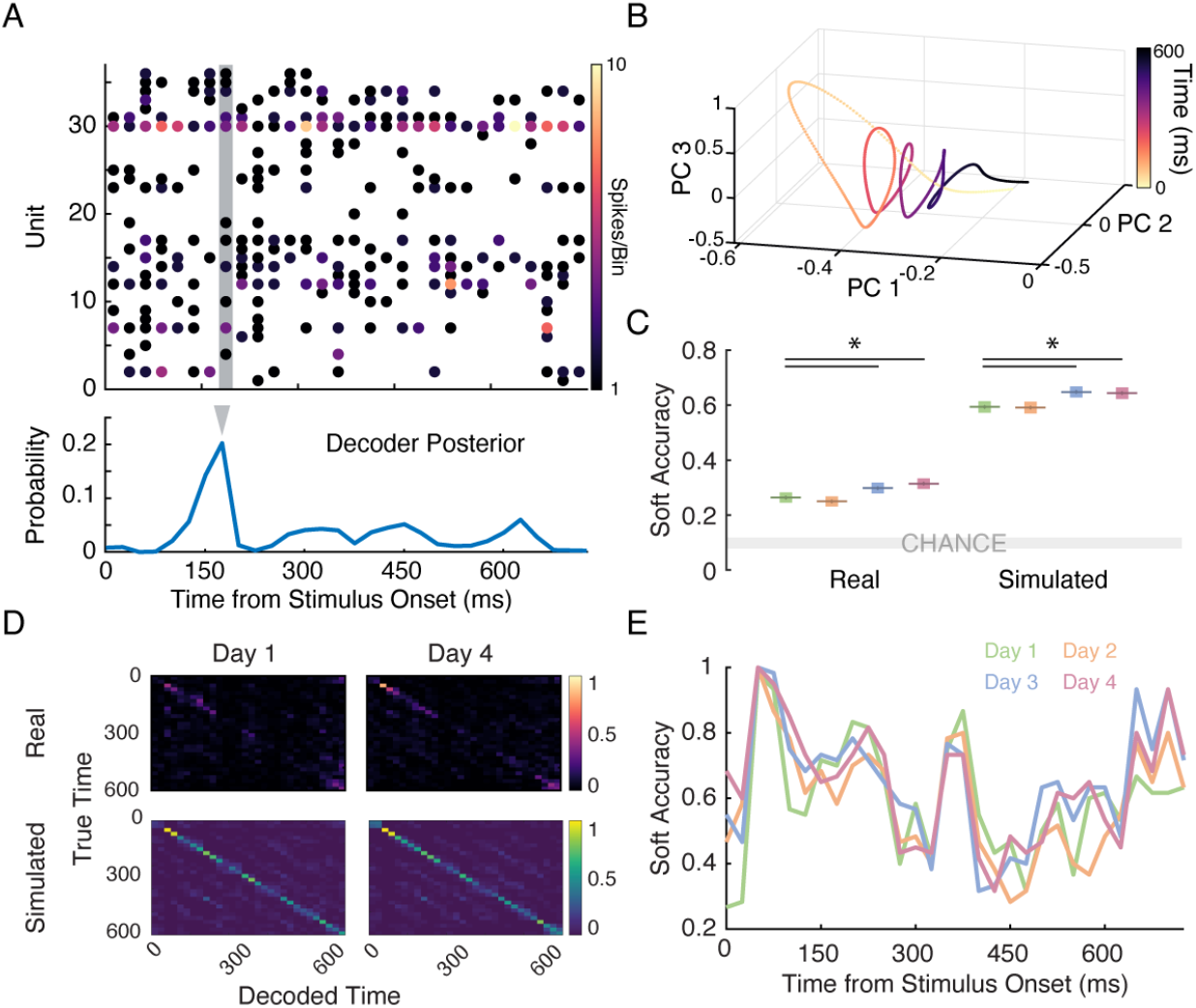
Temporal Information in the V1 Neural Code. **A**. Top: Example pseudo-trial used for instantaneous time decoding. Each row represents a unit (all from the same Test day), while each column represents one 25-ms time bin. Bottom: Posterior probability distribution over possible time bins, calculated for the bin indicated by the shaded box above. In this case, the posterior correctly classifies the time bin based on a maximum a posteriori (MAP) estimate (marked with the gray triangle). **B**. Dynamic latent trajectory representation of ABCD from Day 3-units demonstrate spiraling dynamics (as Figure 3) that create unique locations in PC space for each time point. **C**. Accuracy of temporal decoding across 60 held-out pseudo-trials and 30 time bins / trial for real (left, ∼35 units per day) and simulated (right, 250 units per day, see Methods) data. “Soft accuracy” allows the decoder to be off by one time bin in either direction. Accuracy improved significantly with training (two-sided permutation test for difference in accuracy from Day 1, threshold set to 0.05/3=0.0167 to control for multiple comparisons. Real data: Day 2 p=0.26; **Day 3 p=0.0059**; **Day 4 p<1e-6**. Simulated data: Day 2 p=0.802; **Day 3 p=3.25e-4**; **Day 4 p=6.29e-4**). **D**. Time decoding confusion matrices for naïve (left) and trained (right) units for real (top) and simulated (bottom) data. **E**. Decoding soft accuracy in each time bin for simulated data Training increases accuracy primarily at the beginning and ending of the sequence, which might be explained by the slight increase in evoked firing rate across days of training. Decoding is most accurate around element transitions (0, 150, 300, 450, 600 ms).

As described above, Day 1 has the highest N and model explained variance is no better in trained than naïve mice, so this result cannot be explained by differential fit to the data. However, firing rates did increase across days, which may affect decoder accuracy. Therefore, we removed the highest firing channels, destroying the correlation between firing rate and day (*ρ* = 0.06, p=0.54), then re-ran the decoder. Temporal decoding accuracy still improved significantly between Days 1 and 4: Day 1-22.1%; Day 2-24.1% (p = 0.14); Day 3-21.9% (p = 0.92); Day 4-27.5% (p = 4.6e-5). When we looked more closely at the individual time bins, however, we found that most of the improvement with training occurred during the evoked response to A and the offset at the end of the sequence. We reasoned that temporal decoding performance might be limited by the relatively small number of neurons we recorded from. To investigate further, used MbTDR to simulate a population of 1000 units (250 per day) whose dynamic activity matched the recoded data (Figure 7c-e). The temporal decoder was more accurate with the simulated data (Figure 7c-d) and this accuracy also increased with training, but similar to the real data improved accuracy from Day 1 to 4 was largely restricted to sequence onset and offset (Figure 7d-e). We conclude that evoked neural dynamics in V1 can serve as a reliable source of “when” information for downstream regions and that this information may increase with training. However, our data shows that improvements in decoding accuracy are not uniform across the temporal extent of the sequence, raising the question of whether increases in decoding accuracy provide evidence that the circuits explicitly encode temporal representations. Answering this question will likely require recording cells in other cortical layers and, potentially, other brain regions as well.

## Discussion

We have provided experimental evidence to support several basic principles of predictive processing. We used a statistical model to capture complex and heterogeneous changes in neural population activity across an unsupervised learning process. The model revealed coordinated changes in evoked neural firing that depended strongly on training, including predictive ramping in advance of the expected onset of visual stimuli and spiraling 3D latent trajectories, which might be used as a basis for keeping time. In addition, the model uncovered a significant reduction in firing in a late window after stimulus, from about 100-150ms after both A and B, which progressively decreased with training. Due to this reduction, the unexpected substitution of the trained element A with the novel E caused elevated firing in that same late window. This differential response, consistent with a positive prediction error, was never present in naïve mice. Similarly, the omission of trained element C, which was done by extending the duration of the second element in the sequence, drove sustained, elevated firing in a late window after the expected time of C onset. This result is consistent with a negative prediction error. We note, however, that this was not restricted to sequences obeying the precise trained sequence order, i.e. those starting with AB. Indeed, the negative prediction error was only weakly tuned to the orientation of the preceding elements.

Finally, we used the model to perform instantaneous time decoding, showing that we could reliably decode elapsed time. This result is consistent with V1 being a source of “when” information, as the neural data at different timepoints throughout the sequence was uniquely identifiable. We also note that this ability to decode time is closely related to the concept of temporal redundancy reduction (H. Barlow, 2001a; H. B. Barlow, 1989). Redundancies in a sensory neural code arise when the same sensory information is transmitted at multiple points in space and time. Such redundancies arise from spatial and temporal autocorrelations in the incoming visual signals, which are then propagated by neurons in the visual system. There is significant experimental evidence that the retina and lateral geniculate nucleus (LGN) work to reduce spatial and temporal redundancies in the neural code transmitted to V1 (Dan et al., 1996; Dong & Atick, 1995; Hosoya et al., 2005; Srinivasan et al., 1982). Given the complexity of the incoming signals, however, retinal and LGN processing are likely insufficient to fully remove the autocorrelations and compress the data. V1 might therefore continue the process of redundancy reduction, removing higher-order spatial and temporal correlations, before transmitting to downstream regions. Were this the case, we might expect unsupervised training with a stimulus like ours to reduce temporal redundancies and autocorrelations in the transmitted neural code as the system adapts to the novel stimulus statistics. Though indirect, temporal decoding accuracy might be useful as a surrogate measure for a lack of temporal redundancies.

Three notable recent studies investigated complementary ideas. In the first study, Marina Garrett and colleagues trained mice to perform a visual change detection task in which familiar and novel images were sequentially presented (Garrett et al., 2020). In general, their results closely matched the predictions of efficient coding and predictive processing. Excitatory neuron activity was reduced and more sparsely coded for familiar versus novel images, consistent with the idea that the visual system changed over time to compress its representation of the familiar images. They also showed that L2/3 VIP interneuron activity became predictive of the temporal structure of the sequence. In particular, VIP firing rates ramped in advance of expected stimulus presentations, and ramping activity persisted when an expected stimulus was omitted. In our study, after training, a comparable ramp begins before the onset of each element. In trials where the third sequence element was omitted, the ramp plateaued and sustained for about 100ms until the appearance of the next sequence element. The difference between the continued ramping seen in Garrett et al. and our “ramp-then-sustain” phenotype may be due to recording location and neuron type. In particular, L2/3 VIP neurons that ramp may disinhibit L4 excitatory neurons, whose activity eventually saturates and sustains.

In another study, Peter Finnie et al. analyzed two forms of experience-dependent plasticity in V1 after bilateral lesion of the hippocampus (Finnie et al., 2021). They induced plasticity through sequence learning and also through stimulus-specific response potentiation (SRP). In the SRP protocol, mice passively view a sequence of phase-reversing sinusoidal gratings at 2 Hz (Cooke & Bear, 2010; Cooke, Komorowski, Kaplan, Gavornik, & Bear, 2015; Frenkel et al., 2006). These two forms of plasticity have previously been shown to rely on different mechanisms, and this most recent study further confirmed those mechanistic differences: sequence learning was eliminated in the lesioned animals while SRP was not affected. This suggests that some forms of visual plasticity may require an intact hippocampus and is particularly important given recent investigations revealing strong interactions between the hippocampus and V1 (Diamanti et al., 2021; Fournier et al., 2020). Their study also used visually-evoked potentials to detect the presence of a negative prediction error when the second element was omitted by extending the duration of the first sequence element. Our results replicate theirs with multi-unit data and provide additional insights into potential mechanisms.

In a final study, Colleen Gillon et al. analyzed learning-related changes associated with passive exposure to image sequences (Gillon et al., 2021). They found significant changes in neural activity due to experience, including differences between feedback and feedforward layers of V1, and positive prediction errors in response to unexpected stimuli. In addition, those neurons that responded most strongly to an unexpected stimulus in one imaging session were found to have the largest learning-related changes in subsequent sessions. Thus, unexpected stimuli may preferentially support learning. These results are also consistent with theories of efficient coding and predictive processing. However, it is important to note their conclusions hinged on evidence for a positive prediction error. These can be generated in simple feedforward circuits, for example ones that perform principal components analysis, and are consistent with a wide variety of models of cortical function (Keller & Mrsic-Flogel, 2018). Negative prediction errors, especially those generated in the complete absence of sensory stimulation, are much more difficult to explain and direct evidence for them is minimal [see (Fiser et al., 2016; Keller, Bonhoeffer, & Hübener, 2012)].

To conclude, we would like to emphasize several key points. If the visual system does indeed perform some kind of predictive processing, then its predictions must be based on the environmental statistics that the brain encounters. Therefore, responses to the sequence stimulus in naïve mice, on Day 1, can themselves be thought of as prediction errors to an unexpected stimulus. The sequence stimulus violates known natural environmental statistics and ought to drive plasticity processes in the visual system, which ultimately alter those expected statistics. The evidence collected thus far suggests the mouse visual system does not learn a complete, abstract representation of the sequence, but rather slowly adapts to the new environmental statistics.

In addition, though prediction errors are often emphasized as crucial experimental indicators of predictive processing, they can also be thought of as one example of a more fundamental computational goal, namely information compression (Creutzig & Sprekeler, 2008; Palmer et al., 2015; Srinivasan et al., 1982; Tishby, Pereira, & Bialek, 2000). In this view, incoming sensory data is passed through a bottleneck that reduces the entropy of transmitted data, across both space and time. This is the express purpose of the original predictive coder from telecommunications engineering, which reduces spatial and temporal autocorrelations in the transmitted code (Elias, 1955). Thus, a variety of experimental results, including a reduction in temporal autocorrelations in LGN (Dan et al., 1996) and increasing sparsity in V1 (Failor, Carandini, & Harris, 2021; Olshausen & Field, 1996; Van Vreeswijk, 2001), all point toward some form of efficient coding or predictive processing (Chalk et al., 2018; Niven & Laughlin, 2008; Pitkow & Meister, 2012). A truly efficient code ought to compress information in both space and time. Given our findings, and the convergence of overlapping results in other recent studies, future research might utilize these and other types of sequential stimuli, in conjunction with theoretical tools such as MbTDR or the information bottleneck (Palmer et al., 2015; Tishby et al., 2000), as a reliable way to probe unsupervised learning and sensory processing in the nervous system.

## Methods

### Animals

Male and female C57BL/6 mice (Charles River Laboratories) were housed with same-sex littermates (four mice per cage) on a standard 12-hour light/dark cycle, and provided food and water *ad libitum*. All experiments were performed during the mouse’s light cycle at approximately the same time of day (∼10am-2pm). Mice were 66.8 +/- 0.5 postnatal days old at experimental start (minimum of 61 days and maximum 73 days). Of 56 mice that were considered for further analysis (see Criteria for Multi-Unit Inclusion below), 27 were male and 29 female. There were no significant differences in spontaneous or evoked firing rates between units from male and female mice (Mann-Whitney U test: p=0.17 and p=0.63, respectively). There were also no clear qualitative differences in stimulus-evoked responses, so all analyses were performed ignoring the male/female distinction. All procedures were approved by the Institutional Animal Care and Use Committee (IACUC) of Boston University.

### Experimental Design

Our goal in designing the experiment was to access neural data across the learning process, under the restriction that we could not expect to record from the same units across days. To do so, we created a novel randomized Train/Test protocol (see Figure 1), where mice were selected to see either a Training stimulus or a Test stimulus on any of four experimental days. The randomized selection process acted as a filter, whereby a mouse who saw the Test stimulus was removed from the process and done with the experiment. The overall result of this filter was that ∼25% of mice saw the Test stimulus on Day 1 (the naïve group), ∼25% on Day 2, ∼25% on Day 3, and ∼25% on Day 4. A mouse who saw the Test stimulus on Day 4 would have seen the Training stimulus on Days 1-3, while a mouse who saw the Test stimulus on Day 2 only saw the Training stimulus on Day 1. In order to achieve the desired uniform distribution across days, the following probabilities were used to select mice for Train/Test: Day 1, 75% see the Training stimulus / 25% see the Test stimulus; Day 2, 66.667% Training / 33.333% Test; Day 3, 50% Training / 50% Test; Day 4, 100% Test. Thus, the population of mice remaining in the experiment decreases across days, until they have all seen the Test stimulus. All data analyses were then performed only on neural responses to the Test day stimuli.

### Electrode Implantation

Mice were anesthetized with an intraperitoneal injection of 50 mg per kg ketamine and 10 mg per kg xylazine and prepared for chronic recording as described previously (Cooke & Bear, 2010; Frenkel et al., 2006; Gavornik & Bear, 2014). To facilitate head restraint, a steel headpost was affixed to the skull anterior to bregma using cyanoacrylate glue. Small (<0.5 mm) burr holes were drilled over binocular primary visual cortex (3.1 mm lateral from lambda) and a custom-made recording bundle (20-μm outer diameter tungsten H-Formvar wire, California Fine Wire Company) with 6 wires tightly wound together was placed 450 μm below the cortical surface (Layer 4 / Layer 4/5 border). All recordings were performed in the left hemisphere, as our setup included an infrared camera directed at the right eye. A reference electrode (silver wire, A-M systems) was placed below the dura at ∼1mm lateral from bregma in the same hemisphere as the bundle in V1. All electrodes were rigidly secured to the skull using cyanoacrylate glue. Dental cement was used to enclose exposed skull and electrodes in a protective head cap. Buprenex (0.1 mg per kg) was injected subcutaneously for postoperative pain amelioration. All surgeries were performed around postnatal day 60. Mice were monitored for signs of infection and allowed at least 48 hours of recovery before habituation to the recording and restraint apparatus.

### Data Recording

Each multi-unit bundle contained 4-6 channels per mouse, depending on the custom build process. Each channel was electroplated using a Nano-Z system to yield a final impedance of ∼210+/-10 kOhms at 1000 Hz. Data from each channel was amplified and digitized using an Open Ephys recording system. Spiking activity was digitized at 30-kHz and bandpass filtered from 300-6000 Hz using a causal 2^nd^ order Butterworth filter (implemented in Open Ephys). This bandpass data was extracted from a binary storage format and analyzed using custom software written in MathWorks MATLAB (see below). Mice were head-fixed and awake during all recordings.

### Criteria for Multi-Unit Inclusion

The bandpass data from a single mouse and session, ***Z*** ∈ ℝ^(*recording time*)*x*(*channels*)^, was first transformed to its singular value decomposition (***Z*** = ***USV***^*T*^), yielding an orthogonal set, ***U***, and placing movement artifacts into a single dimension (invariably the first dimension, with the most variance). Each column of the matrix ***U*** was then reduced to a set of timestamps representing multi-unit spike times. This was done by identifying all those times the voltage trace crossed a negative-going threshold of 4 robust standard deviations from the mean (the robust standard deviation was estimated as 1.4826 times the median absolute deviation). A given effective channel (column of ***U***) was then selected for further analysis if it met the following criteria:

1. Had a mean evoked firing rate greater than 1 Hz (with a sequence of 4 elements and 150ms / element, 1 Hz is less than 1 spike per trial)
2. Passed a statistical test for visual responsiveness
3. Had no more than R=0.258 stimulus-evoked correlation with all other simultaneously recorded channels (R=0.258 is equivalent to R^2^=0.067, which was our estimate of the proportion of explainable variance in our experimental setup)
4. Showed no signs of artifacts in spike raster

The test for visual responsiveness consisted of two statistical tests, and each effective channel had to pass at least one of the tests with a Bonferroni-corrected p-value <= 0.01/2. The visual stimulus had four visual elements and a fifth “element” driven by the offset response as the screen switched back to gray. The first test was a two-sided KS test, which compared the evoked distribution of inter-spike intervals (ISIs) in a window spanning all five visual elements to a null distribution of comparable size drawn from periods with no visual stimulation. This test captures multi-unit channels with evoked distributions unique from their spontaneous distributions. The second test was a chi-square difference of deviance test. For each effective channel, we fit two Poisson GLMs to model the evoked peri-stimulus time histogram (PSTH). The first model, null, had 2 parameters such that the evoked PSTH was restricted to be a linear function of the time elapsed in the sequence. The second model, full, had 1+T parameters for 1 baseline parameter and T time bins (T=30 for 25ms bins and 150ms*5=750ms total time in a sequence). We then compared the difference of their deviances to a chi-square CDF with degrees of freedom given by the difference between the number of parameters in the two models (T-1). This test captures multi-unit channels with “interesting” PSTHs (those with more wiggles than a linear function). The final result of this process preserved 140 distinct multi-unit channels (of a total 347 potential channels) from 56 mice (of 67 mice implanted).

### Visual Stimulus Presentation

Visual stimuli were generated using custom software written in Matlab with the PsychToolbox extension (http://psychtoolbox.org/) to control stimulus drawing, timing, and synchronization with the Open Ephys. Stimuli were presented on a screen placed 25cm from the mouse, directly in front to target stimulation of binocular V1. The stimulus consisted of a sequence of 4 visual elements, followed by an inter-sequence gray period. Each visual element persisted on screen for 150ms, while the gray period was randomized from a uniform distribution between 1 and 2 seconds. The total duration of a sequence was 4*150ms=600ms. Each visual element was a full-screen oriented sinusoidal grating (0.05 cycles per degree), shown at 75% contrast. For consistency, we used the same orientations across mice: A-75°, B-120°, C-35°, D-160°, E-10°, *x*-Uniform (60°,180°). Grating stimuli were gamma corrected to insure a linear gradient and constant total luminance. Stimuli were also corrected to ensure the gratings had a constant spatial frequency in retinotopic coordinates. This consisted of a spherical transformation, as described in (Marshel, Garrett, Nauhaus, & Callaway, 2011), supplemental information. During experiments, animal handling consisted of placing each mouse into the head-fixed presentation apparatus. The handler was unaware of whether the mouse would see the Test or Training stimulus.

Training Stimulus – If a mouse was randomly selected to see the Training stimulus, it saw 200 presentations of the sequence ABCD. Each sequence was presented in four blocks of 50 presentations, with each presentation separated by a gray screen drawn from a randomly-selected uniform interval of 1 to 2 seconds, and each block separated by 60 seconds. The total time of visual stimulation was 200*600ms = 2 minutes and the total time in the apparatus ∼10 minutes.

Test Stimulus – If a mouse was randomly selected to see the Test stimulus, it saw 600 presentations of sequences from one of four conditions: A*x*CD, A*x*→D, E*x*CD, E*x*→D. *x* is a randomly oriented element and *x*→ means the second element was held on screen for 300ms, such that nothing changed at the B-C transition time observed during Training. As before, each sequence was presented in blocks of 50 presentations, but now for a total of 12 blocks. The order of the presentations and the orientation of the second element, *x*, was randomized for each Test Day, i.e. all mice seeing the Test stimulus on Day 1 saw the same set of sequences in the same order, while mice on Day 2 saw a different set in a different order (this allowed for the creation of pseudo-trials with data grouped across all mice recorded on the same Test Day). The total time of visual stimulation was 600*600ms = 6 minutes and the total time in the apparatus ∼30 minutes.

### Data Analysis

All analyses aside from the explainable variance estimation were performed only on Test day data, so there were no repeated measures. In general, we had N=140 multi-unit channels from the Test days, with 37 units on Day 1, 31 on Day 2, 36 on Day 3, and 36 on Day 4. Therefore, if we performed a statistical test comparing Day 1 and Day 4, there would be 37+36=73 data points, assumed independent and identically distributed within a test day group as most units came from different mice and those that came from the same mouse were required to have a very low pairwise evoked correlation (see Criteria for Multi-Unit Inclusion above). Any figure depicting a bootstrap confidence interval was computed as a bootstrap pivotal interval (Wasserman, 2004).

### Pre-Processing

Firing rate was first computed as a spike count in 25-ms bins, with no smoothing. Each unit’s firing rate was then normalized by subtracting out the spontaneous baseline firing rate and dividing by the evoked standard deviation to yield zero baseline mean and unit variance. This z-scored firing rate was used in all analyses and is displayed in all figures (the notation *FR (AU)* used in multiple figures refers to this z-scored firing rate, while the notation *FR Modulation* is reserved for shared bases computed from MbTDR, see below). Dot plots (as in Figure 3b) were computed directly from the z-scored firing rates, averaged in 50-ms windows as described in the main text. In the case of the firing rate difference plots (Figure 4c-d and Figure 5c-d), we first computed for each unit separately z-scored PSTHs for the two stimuli being compared (e.g. A/E*x*CD and A/E*x*→D). Next, we subtracted one PSTH from the other to create a difference PSTH. Finally, we averaged the difference PSTH in a 50-ms window to yield a *Mean FR Difference*. Statistical tests were then performed on the FR differences across Test days (see below).

### Explainable Variance Estimation

In order to estimate the expected proportion of variance explained for our setup and stimulus, we utilized data recorded on the Training days. On those days, each animal saw 200 presentations of the same stimulus, ABCD. Therefore, the PSTH provides an accurate estimate of the neural variability explainable by the stimulus. Using the same criteria for multi-unit inclusion as used on the Test days, we compiled a set of 156 unique multi-unit channels. For each unit, we fit a Poisson GLM to model the PSTH (as in the Criteria for Multi-Unit Inclusion section, with 1+T parameters for 25ms time bins and T=30). We then calculated the proportion of neural variance explained by the model:

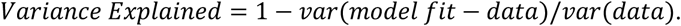

The average variance explained from these 156 channels provided our estimate of explainable variance: 6.7 +/- 0.7% (mean +/- SEM). Thus, a substantial proportion of the neural variability cannot be captured by a model that simply accounts for the visual stimulus, as anticipated by other research on the mouse visual system (Stringer, Pachitariu, Steinmetz, Carandini, & Harris, 2019; Stringer et al., 2018).

### Model-Based Targeted Dimensionality Reduction (MbTDR)

Many of the data analyses were performed by fitting a single supervised statistical model to all recorded multi-unit data across the four Test days. The model is known as model-based targeted dimensionality reduction (Aoi et al., 2020; Aoi & Pillow, 2018), which reduces dimensionality by projecting the data onto low-dimensional subspaces corresponding to different covariates. The covariates in this case are those related to the structure of the experimental design and the Test day stimulus set: condition-independent (a constant value of 1), experimental day, element A or E to start the sequence, the orientation of the second element (*Angle*(*x*)), whether the second element was held on screen [*x*→], trial number, and interaction terms between these. We also included several triplet interaction terms: *Angle*(*x*) * *I*(*E*) * *Day, Angle*(*x*) * *I*(*x*→) * *Day, Angle*(*x*)^2^ * *I*(*x*→) * *Day, Angle*(*x*)^2^ * *I*(*E*) * *Day*, none of which were ultimately chosen in the model selection process. Note that all covariates were designed so that the baseline corresponds to the sequence ABCD on Day 1. For example, day is encoded by log(*Day*) so that the covariate for Day 1 is zero, A or E to start the sequence is encoded as an indicator function *I*(*E*), and the orientation of the second element, *x*, is centered so that the orientation of B used during training (120°) is equal to zero. For more information on these covariates and how they were encoded, see Supplemental Table 1. The output of the model is comparable to de-mixed principal components analysis (dPCA) (Kobak et al., 2016).

We will now describe the MbTDR model and how it derives from a standard linear regression (for a more complete description, see (Aoi & Pillow, 2018)). For the present dataset, we recorded from *N* = 140 multi-unit channels and *M* = 600 trials/unit. The data was binned at 25ms, with *T* = 30 time bins per trial for a total trial time of 750ms (4*150ms for each of the visual elements and 150ms for the offset response as the screen turns back to gray). Each unit’s firing rate was first z-scored as described above.

Assuming the covariates vary by trial only (not within a trial) and we wish to infer how neural activity evolves over time on individual trials, we might create a linear model for one unit, *n*, on one trial, *m*:

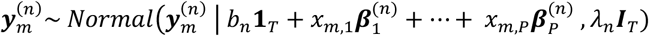

where 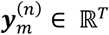 is the firing rate of unit *n* on all *T* timepoints of trial *m*, each 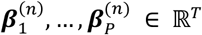 is a regression coefficient vector for unit *n* that covers the *T* timepoints of a single trial, *b*_n_ is a unit-specific baseline firing rate, *λ*_*n*_ is a unit-specific variance, and *x*_*m,p*_ ∈ ℝ is the value of the *p*-th covariate on trial *m*. Each column of 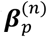 is therefore the PSTH of the neuron projected onto the subspace spanned by the *p*-th covariate. If we assume the baseline firing, *b*_*n*_, variance, *λ*_*n*_, and covariate-specific bases, 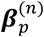, are constant across trials, then we could use a least-squares approach to learn the parameters.

Recording from *N* units, with *T* timepoints per trial, and using this linear model, we would find a set of parameters, 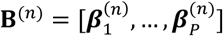, for each unit, plus the baseline firing rates and noise variances. For the entire dataset, that is 2*N* + *TPN* parameters (∼140,000 in the present case). With the limitations on recording time in animal experiments, and the variability of neural data, collecting enough trials to reliably estimate so many parameters would be quite difficult for more than a few covariates.

The key idea of MbTDR is to reduce the total number of parameters by attempting to discover a low-rank representation for the regression coefficients corresponding to each covariate. That is, find 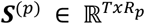 and 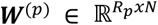 such that:

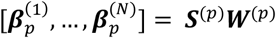

where *R*_*p*_ is the rank of the representation for covariate *p*. This low-rank representation effectively discovers a unique set of shared time-varying basis functions, ***S***^(*p*)^, for each covariate, which are modulated independently for each unit by the latent factors, 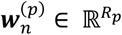. The shared basis functions are a low-dimensional representation of high-dimensional neural trajectories. With this adjustment, the total number of parameters can be dramatically reduced to 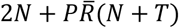), where 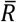 is the average rank (∼4,500 in the present case).

Under MbTDR, the full likelihood of all recorded data is:

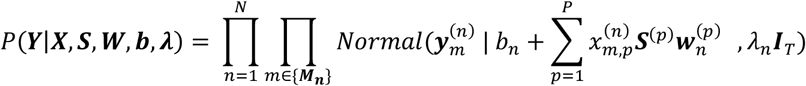

where now ***Y*** is the set of all 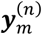, ***X*** is the set of all trial covariates, ***S*** and ***W*** are the sets of all ***S***^(*p*)^ and ***W***^(*p*)^, ***b*** is the vector of all *b*_*n*_, ***λ*** is the vector of all *λ*_*n*_, and ***M***_***n***_ is the set of all trials unit *n* participated in.

To fit this model, the original authors proposed an expectation conditional maximization either algorithm (ECME), so the optimization is actually over a marginal likelihood: *P*(***Y***|***X, S, b, λ***). We wrote custom scripts to perform ECME for initialization and then passed the result to a trust-region algorithm (Matlab *fminunc*) to directly maximize the marginal likelihood. To compute the marginal likelihood, all of the “unit factors” are given a *Normal*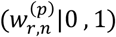 prior and then integrated out. This conjugate Gaussian prior, along with the independence of the unit factors, dramatically decreases the complexity of both the derivation and the optimization problem. The optimal set of parameters are then maximum marginal likelihood estimates 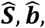, *and* 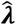, along with the posterior mean for ***W***, computed from 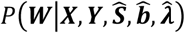.

To determine the optimal rank of the coefficient matrix for each covariate, we performed a greedy forward stepwise algorithm that sought to minimize the Akaike Information Criterion (AIC) (again following (Aoi & Pillow, 2018)). To begin, we fit the model independently *P* times (with the *p*-th covariate set to rank 1 and all others to 0), and then chose the covariate for which the AIC was minimized, *p*_*min*_ [1]. With the rank of covariate *p*_*min*_ [1] now equal to 1, and all others still set to zero, we repeated the process by again fitting the model *P* times, iteratively adding 1 to the rank of each covariate, and finding the minimum AIC across the *P* iterations, *p*_*min*_[2]. Now, with *p*_*min*_ [1] set to 1 and *p*_*min*_ [2] set to 1, the process continued (if *p*_*min*_ [1]= *p*_*min*_ [2], then that covariate would be at 2 with all others still equal to 0). This was repeated until the global minimum AIC was achieved. 90% of the data (540 trials / unit) was used for this fitting procedure. The remaining 10% (60 trials / unit) was excluded from all preliminary analysis and model fitting, and reserved for decoding and for estimation of the held-out proportion of variance explained by the model.

With the optimal model parameter estimates, we can evaluate the model fit for each unit and stimulus type:

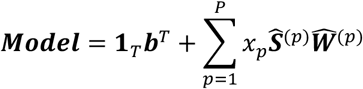

Where **1**_*T*_ is a vector of *T* ones, *x*_*p*_ is a scalar representing the value of the *p*-th covariate for the desired trial type, and ***Model*** ∈ ℝ^*TxN*^ are the model fits. To evaluate the model response to the training sequence ABCD, we simply set the covariates appropriately (i.e. *I*(*x*→) = 0, *I*(*E*) = 0, *Angle*(*x*) = 0). To determine the orientation tuning of each unit to the second element when A starts the sequence, we can set *I*(*x*→) = 0 and *I*(*E*) = 0, and then change the value of *Angle*(*x*) across its range from 60 to 180 degrees. This yields a PSTH for each unit and each angle, from which we then extract the maximal response in a window from 50-150ms after the onset of the second element (200-300ms after the onset of the entire sequence). The set of maximal responses across possible angles is the orientation tuning curve for one unit. Taking the angle for which the orientation tuning curve is a maximum gives the peak tuning angle for that unit.

For visualization purposes and to compare the low-rank representation across conditions, as in Figures 3-5, we performed the following singular-value decomposition:

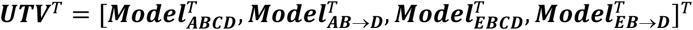

Thus, ***U*** ∈ ℝ^*TTx*(*total rank*)^, and every column of ***U*** represents a set of low-dimensional neural trajectories that can be reliably compared across conditions (the notation FR Modulation, used on the y-axis of multiple figures, refers to the columns of ***U***, which are normalized by the SVD procedure). The first *T* elements of the first column of ***U*** are the response to *ABCD* in the dimension with the greatest singular value, while the next *T* are the response to *AB*→*D* in that same dimension. To make comparisons across days (as in Figure 3c-f), we restricted the set of units used in the model fit such that, for example, 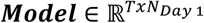. Next, we computed the singular-value decomposition with that restricted set of model PSTHs, and then repeated for each Test day.

The variance accounted for by each component, *i*, is computed with ***T***:

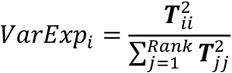

This is the variance explained by each component in the singular-value decomposition of the model fits: it is variability in the PSTHs rather than the neural data, so it represents signal rather than noise variance.

### Decoding Pseudo-Trial Covariates

In Figure 2, we showed several examples of stimulus decoding. MbTDR allows for decoding by invoking Bayes’ rule:

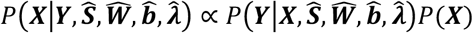

In our experiment, all covariates were randomly generated from uniform distributions, so *P*(***X***) is a constant that can be ignored. Even though the base model is linear in its covariates, the optimal decoder is non-linear, so we again used a trust-region algorithm to maximize 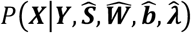 with respect to the covariates ***X*** on individual pseudo-trials. In particular, the set of covariates are created from a “fundamental” set, e.g. *Angle*(*x*)^2^, and *I*(*x*→) * *Day* come from the more fundamental *Angle*(*x*), *Day*, and *I*(*x*→). A linear decoder would provide different estimates for each covariate, e.g. 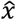 and 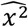 such that 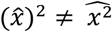. The non-linear decoder, however, outputs point estimates of the fundamental set of covariates, given the neural data and the optimal model. Those estimates include: experimental day, angle of the second sequence element (*Angle*(*x*)), trial number, a posterior probability for the indicator of E starting the sequence (𝔼[*I*(*E*)]), and a probability for the indicator of the third element having been omitted (𝔼[*I*(*x*→)]).

Pseudo-trials are not simultaneously recorded but are created by grouping together single-trial data across all neurons recorded on the same Test day. For example, 37 units from 15 mice were recorded on Day 1. Those mice all saw the same stimuli in the same order, so each trial is comparable across mice. To create a Day 1 pseudo-trial, we selected one of the held-out trials and created a joint log likelihood during that trial by summing over all 37 units. We then found the set of fundamental covariates, 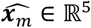, that maximized the joint log likelihood across those units for a given held-out trial, *m*:

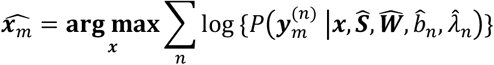

### Decoding Time

In Figure 7, we performed time decoding. Data from each 750ms trial was binned at 25ms, creating a total of 30 bins per trial. The problem of decoding time on a single trial is equivalent to asking which bin from the model fit is most consistent with the data. We can create a scalar normalized firing rate for one unit, timepoint, and trial: 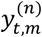. Then, the likelihood of the data in that one time bin is:

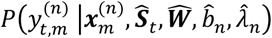

We take the model fit, 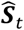, at the same bin and have assumed that the trial covariates, 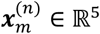, are their true values, i.e. those observed by the mouse on that trial.

If we were to decode time with the data from one unit, the posterior distribution over potential model time bins, *τ*, for a specific bin of neural data, *t*, would be:

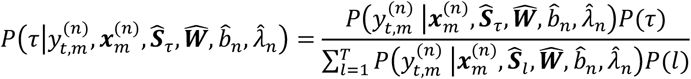

Because the prior distribution over time bins, *P*(*τ*), is uniform, the optimal decoded bin considering all units recorded on a given Test day, trial, and true time bin (*t*) is:

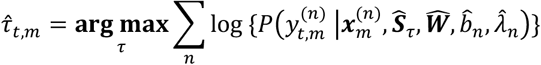

where we vary *τ* from 1 to *T*, selecting the optimal time bin as the one that maximizes this joint log likelihood. This is the time decoder we used in the Results section, which might be called a “conditional” decoder because the time-bin posterior is conditional on the true values of the covariates, 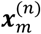. To quantify the accuracy of this decoder, we computed a “soft accuracy” on the decoded time bin for each day:

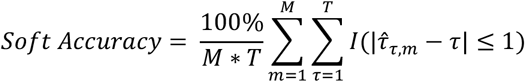

The soft accuracy allows the decoder be off by one time bin in either direction, so there is a 3/30 chance of guessing correctly for the central bins (bins 2-29) and a 2/30 chance on the ends (bin 1 and bin 30), which averages to 9.77%. A 95% confidence interval on chance soft accuracy (with *M* = 60 trials and *T* = 30 time bins on a given Test day) is [8.4, 11.3]%. Therefore, on a given day, a soft accuracy greater than about 12% performs significantly better than chance. In order to determine whether decoding accuracy improved significantly with Training, we performed a permutation test described in the next section. Assuming only one time bin, the 95% confidence interval is [2.1, 17.5]% for 60 trials.

To test the robustness of the result, and in order to more accurately capture the uncertainty faced by the nervous system, we also tried a “marginal” decoder that integrates out the trial covariates. For one unit:

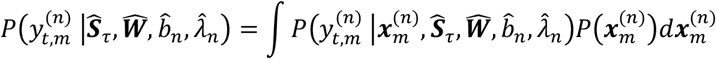

To compute the integral across all units, we used a Monte Carlo sampling procedure. First, we randomly drew a set of trial covariates, ***x***^(*iter*)^, from their respective *a priori* uniform distributions:

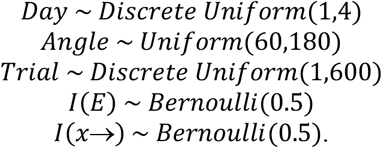

Next, we computed the joint log likelihood of the data from one time bin given those sampled covariates:

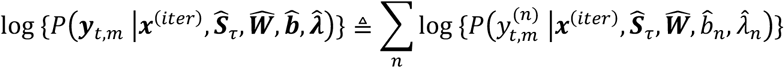

Then, we repeated this process for each of the *τ* = 1, …, *T* bins of the model, ultimately computing a conditional posterior distribution over bins:

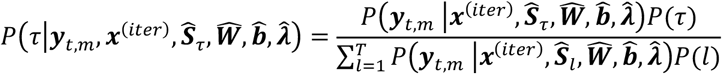

Finally, we sampled from this distribution (which is multinomial), yielding one sample from the marginal posterior:

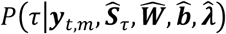

Repeating this process for 1000 iterations, the optimal time bin is then the *τ* with the greatest number of posterior samples. The results were comparable to those with the conditional decoder, though accuracy dropped somewhat. Bootstrap 95% confidence intervals on the soft accuracy were: Day 1- [22.0%,25.6%]; Day 2- [21.6%,25.1%] (p = 0.72, two-sided difference in soft accuracy from Day 1 permutation test); Day 3-[24.8%,28.4%] (p = 0.037); Day 4-[26.1%,29.8%] (p = 2.5e-3).

Using MbTDR, we also simulated data from 1000 neurons (250 per day), matching Test days for noise variance. To simulate data, we discovered kernel-density estimates for the distributions of unit factors (***W***) unique to each Test day (see, for example, Supplemental Figure 3a), and for the noise variance (***λ***) irrespective of Test day. Next, we randomly drew from those distributions, creating new units for each Test day whose values for the unit factors matched the corresponding Test day’s actual neural data. With 60 trials of simulated data, we then performed time decoding exactly as we did for the real data. With more units, decoding soft accuracy improved significantly to ∼60% (up from ∼27% in the neural data, where we had ∼35 neurons per Test day): Day 1-59.4%; Day 2-59.1%; Day 3-64.7%; Day 4-64.4%

### Non-Parametric Statistical Tests

In the Results section, we performed several non-parametric permutation tests to determine the significance of median differences, mean differences, and correlations between evoked firing rates, firing rates & held-out explained variance, decoded covariates & ground-truth covariates, etc. In each case, the null distribution of our test statistic may not follow a known distribution, so we resorted to non-parametric tests (controlling for multiple comparisons when applicable). For the correlation, we performed an approximate two-sided permutation test using Spearman’s rho as the test statistic. For the example of firing rate and explained variance, we have a dataset: 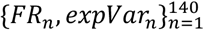 for the complete set of multi-unit channels. We first calculated the observed test statistic by Spearman’s rank correlation: *ρ*_*obs*_. Next, we permuted the dataset so as to break dependence between the two variables, e.g. now the firing rate from the 1^st^ unit is associated with the explained variance of the 17^th^ unit. Then, we calculated a permuted correlation: *ρ*_*perm*_. We repeated this process *L* times by randomly permuting the dataset and calculating new values for 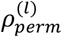. The p-value is then:

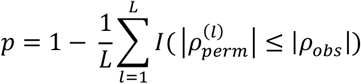

In every case, we performed *L* = 1*e*6 permutations, such that an estimated p-value of zero implies the true p-value is less than 1e-6.

The permutation test on the decoder soft accuracies (Figure 7c) was performed as follows. For each Test day, we had 60 trials and 30 time bins per trial to decode. Thus, the decoder output on a given Test day can be written as a matrix, 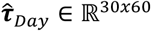. After computing the soft accuracy from this matrix, we calculated a difference in soft accuracy from Day 1 statistic:

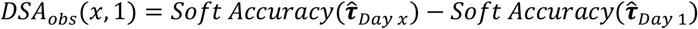

Next, we computed a null distribution for this statistic using random permutations of the data. The null distribution is the distribution of soft accuracy differences, were the soft accuracy the same on both days. For example, if we have decoder outputs from Day 1 and Day 3, 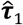 and 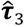, then the full dataset can be written: 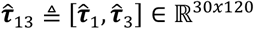. The permuted dataset randomly shuffles values in each row of this matrix (preserving the time bin information), and then re-isolates decoder output matrices to compute the permuted soft accuracy difference:

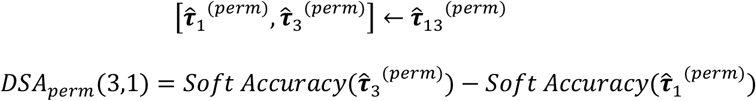

The test’s p-value is then:

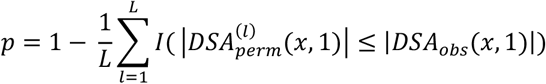

A similar procedure was used in all of the permutation tests from Figures 3-5, with *L* = 1*e*6 permutations.

See Supplemental Table 2 for a summary of all the statistical tests used, along with summary statistics of the data.

## Acknowledgements

Thanks to Mikio Aoi for discussions about the statistical model, MbTDR. Also thanks to Scott Knudstrupp and other members of the Gavornik lab for feedback throughout the process. This work was supported by NEI R01EY030200.

## Author Contributions

Experimental conception and design: JPG and BHP. Experiments and surgeries: BHP, CMJ, AAK. Data analysis and figures: BHP, CMJ, JPG. Writing: BHP, JPG, CMJ.

## Notes

### Competing Interest Statement

The authors have declared no competing interest.

