## Supplemental Figures and Tables for "Expectation violations produce error signals in mouse V1"

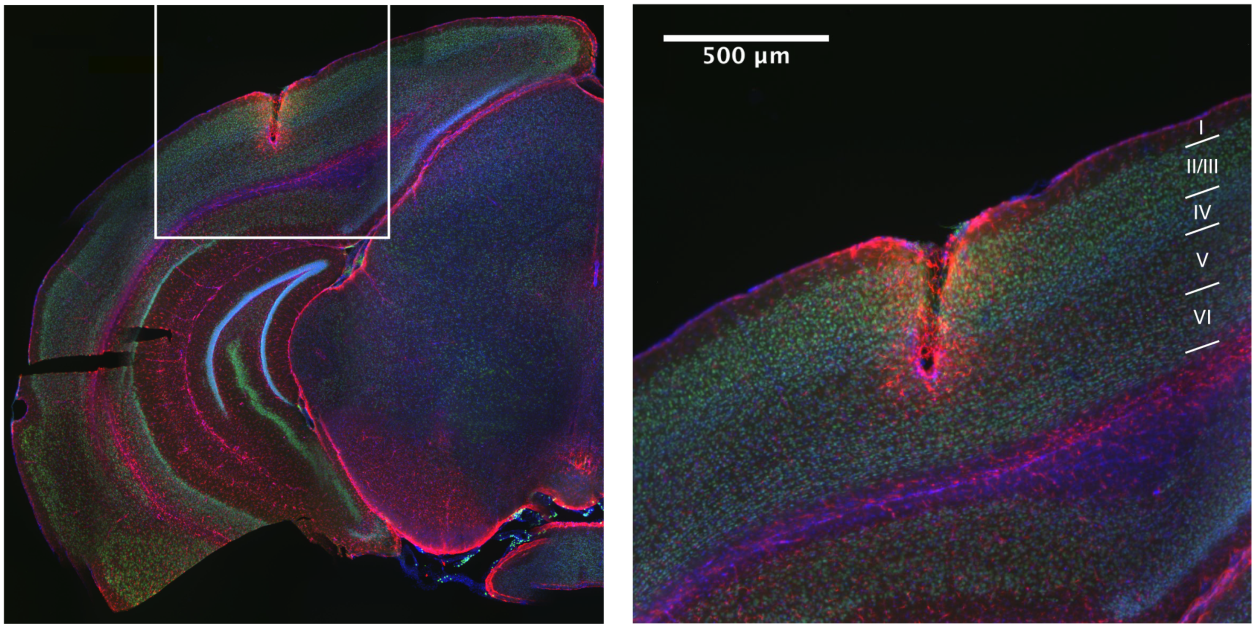


**Supplemental Figure 1: Post-Mortem Histology**
Immunofluorescence image of 50-micron thick coronal section through V1 showing representative electrode placement in binocular Layer 4, near Layer 4/5 border. Blue labeling represents nuclei stained by Hoechst, green represents neuronal nuclei stained by NeuN, and red represents astrocytes stained by GFAP (glial fibrillary acidic protein).

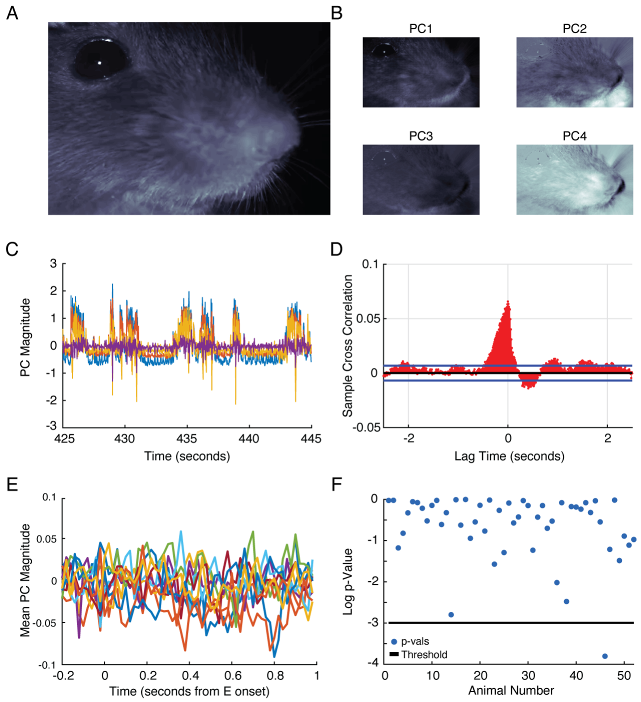


**Supplemental Figure 2: Absence of Stimulus-Aligned Movement**
**A.** Example image of the right eye and whisker pad of a mouse in our apparatus captured at 50Hz using an infrared camera. The cheek and nose move in and out of focus depending on the mouse's facial position. **B.** First four principal component (PC) eigenvectors of a motion energy movie of the mouse's face. These were produced by performing an online Expectation Maximization (EM) algorithm on the absolute value of the difference between adjacent movie frames. Video was acquired during every recording session for each mouse. The first PC represents global movement of the snout and whisker pad, while the second seems to capture snout movements alone. The third and fourth seem to capture more precise movements of the whisker pad, in opposing directions. Other PCs are less identifiable. **C.** Example traces of the first four principal components from a randomly selected 20-second time window. Periods of general quiescence are punctuated by robust movement epochs. Rhythmic activity during quiescent periods is due to a periodic sniffing behavior. **D.** Sample cross-correlation between the first motion energy PC and the recorded neural data from one visually-responsive multi-unit channel. Blue lines show a 95% confidence interval of the null cross-correlation. Negative lags indicate neural data leads the movement signal. **E.** The mean of each of the first ten PCs, from one mouse, aligned to the onset of the deviant stimuli E$x$ →D or E$x$CD. Note the PCs were normalized to have unit variance (see y-axis scale in C), so fluctuations from zero are quite small here. **F.** p-values from likelihood ratio tests for deviant-stimulus-aligned movement in the first PC. For each mouse that had movement and co-recorded neural data from visually-responsive units (52 out of 56 animals), we fit two statistical models that captured the deviant-stimulus-aligned movement (modeling the dark blue trace in E) from the first PC. The full model used a set of Gaussian radial basis functions to capture the stimulus-evoked movement assuming normal residuals, while the null model assumed there was no stimulus-evoked movement. 51/52 (98.1%) of the tests failed to reject the null (all those dots above the black p-value threshold line of 0.05). As a control comparison, we randomly shifted the movement data (preserving its statistical structure but decoupling it from the timing of the stimulus), and found 52/52 tests failed to reject the null. Thus, there was no evidence for deviant-stimulus-aligned movement. In the one case with a p-value less than 0.05, the effect size was small (maximum average stimulus-aligned movement about 1/10 of 1 standard deviation). We found a comparable effect when looking at all stimulus-aligned movement (rather than restricting the analysis to deviant stimuli). While movement provides a meaningful explanation of neural variability generally, the weight of the evidence indicated that the mouse's movement was not aligned to the timing of visual stimulation and therefore its effect would average out in any analysis that was locked to stimulus timing. We therefore did not include movement data in our analyses.

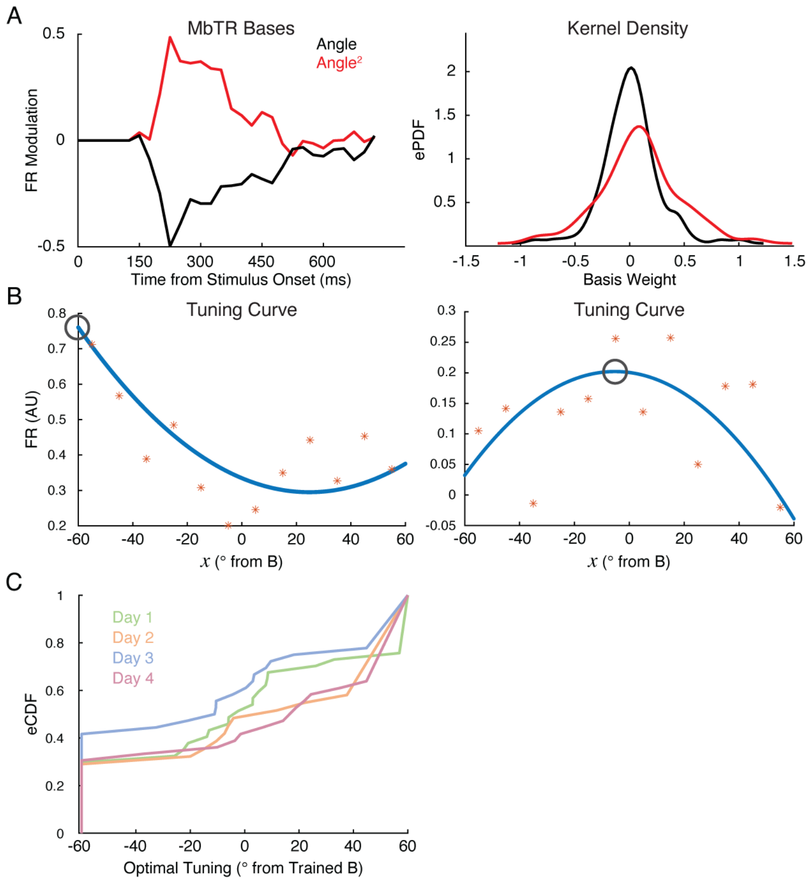


**Supplemental Figure 3: Orientation Tuning of the Second Sequence Element**
**A.** Left: shared basis functions for the $Angle(x)$ (black) and $Angle(x)^2$ covariates (red). Right: kernel density estimates of the unit factor distribution for the same covariates. These bases, along with the rest of the model, can be used to create a tuning curve for each unit at different timepoints after stimulus onset. **B.** Example tuning curves (blue) for two units, taken at the time bin with maximal evoked firing (50-75 ms after the onset of the second element, i.e. the peak in A). Red stars are the raw data, averaged in bins of 10 degrees. The optimal tuning for each unit is the maximum of these curves (black circle). **C.** Empirical cumulative distribution functions for optimal tuning on each Test day. These eCDFs were *not* significantly different between naïve and trained groups (two-sided KS test, naïve [Day 1] vs. trained [Days 3 & 4]: D=0.106, p=0.932).

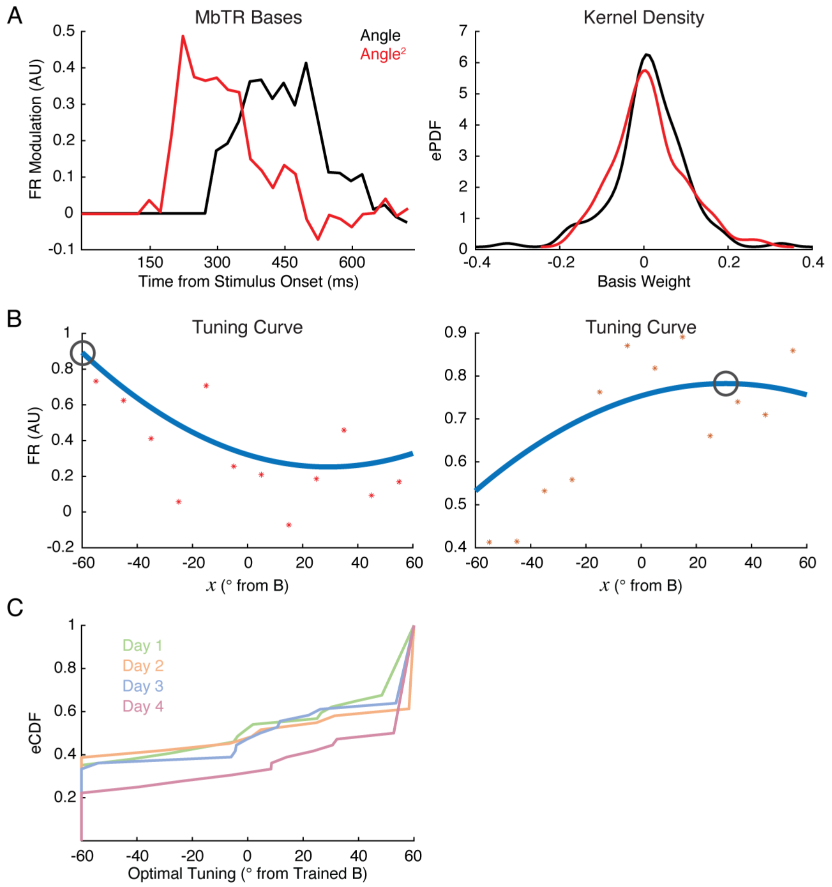


**Supplemental Figure 4: Orientation Tuning of the Negative Prediction Errors**
**A.** Left: shared basis functions for the $I(x\rightarrow) * Angle(x)$ (black) and $Angle(x)^2$ covariates (red). Note their significant overlap in time. Right: distribution of unit factors for the same covariates. Comparable to Supplemental Figure 3, we created negative-prediction-error orientation tuning curves from these bases for trials when the third element was omitted: $I(x\rightarrow) = 1$. **B.** Example tuning curves (blue) for two example units, taken in the late window after the expected onset of C (401-450ms after sequence onset). Red stars are the raw data, averaged in bins of 10 degrees. The optimal tuning for each unit is the maximum of these curves (black circle). **C.** Empirical cumulative distribution functions for optimal tuning on each Test day. These eCDFs were *not* significantly different between naïve and trained groups (two-sided KS test, naïve [Day 1] vs. trained [Days 3 & 4]: D=0.152, p=0.592). A very similar result holds for the early window (D=0.139, p=0.697).

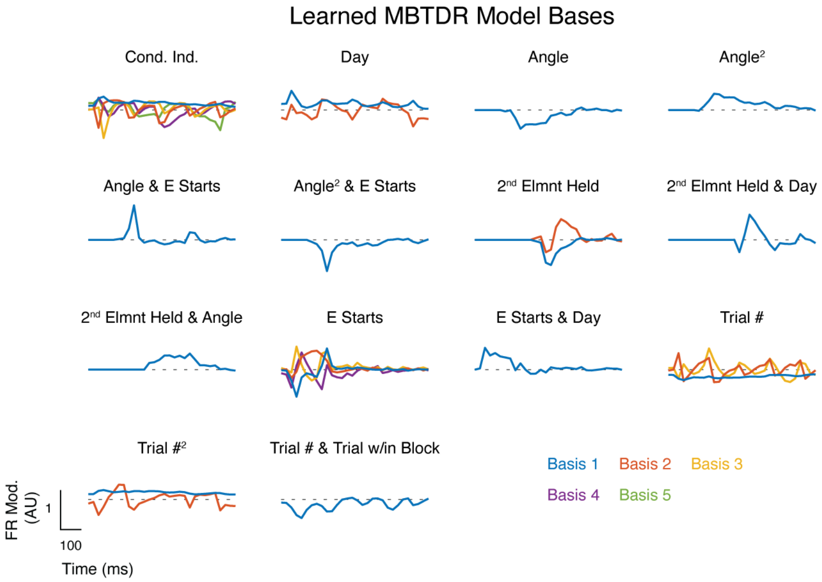


**Supplemental Figure 5: MbTDR Bases**

Visualization of the learned bases ($\boldsymbol{S}$) from the MbTDR fit. All covariates are represented in the model by a matrix $\boldsymbol{SW}^T$, from which we extract a singular-value decomposition $\boldsymbol{UTV}^T = \boldsymbol{SW}^T$. Depicted are the $\boldsymbol{U}$ for each covariate. The bases are all aligned so that $median(\boldsymbol{V}_i) > 0$ for each column ($i$) of the basis; upward deflections of the depicted basis therefore imply a positive change in firing rate for the majority of units. From top to bottom and left to right, the bases are: 1,1: condition independent; 1,2: Day; 1,3: Angle; 1,4: Angle Squared; 2,1: Angle & E starts; 2,2: Angle Squared & E starts; 2,3: Second Element Held; 2,4: Second Element Held & Day; 3,1: Second Element Held & Angle; 3,2: E Starts; 3,3: E Starts & Day; 3,4: Trial Number; 4,1: Trial Number Squared; 4,2: Trial Number & Trial Within Block Number.

**Supplemental Table 1: MbTDR Information**

| Fundamental Covariate | Encoding | Start Time | Final Rank |
|---|---|---|---|
| Condition Independent | 1 | 0ms (beginning of the sequence) | 5 |
| Day | $\log(Day)$ | 0ms | 2 |
| Angle | $Angle(x) - Angle(B)\ radians$ | 150ms (onset of second element, $x$) | 1 |
| Second Element Held | $I(x \rightarrow)$ | 300ms (expected onset of third element, C) | 2 |
| E starts | $I(E)$ | 0ms | 4 |
| Trial Number | $-\log(trial\ \#)$ | 0ms | 3 |
| Trial within Block Number | $-\log(trial\ \#\ in\ block)$ | 0ms | 0 |

Note that $I(*)$ is the indicator function. Each basis spanned the time of the trial (750ms, or 30 time bins), except for the angle and second-element-held bases. For those, we forced the basis to be zero at the beginning of the trial to avoid overfitting (the second element, for example, does not come on screen until 150ms into the trial, so its angle could not possibly have an effect until that moment). $Day$ was simply the test day, from 1 to 4. The logarithm causes this covariate to be zero on Day 1 and to grow sub-linearly beyond that (as we have observed in previous studies with this sequence protocol). Note that because the same units were never recorded across days, this $Day$ covariate acts like an indicator function for training. If some underlying mode of neural activity was common across both naïve and trained mice, it would be picked up by the condition independent component. Alternatively, since the $Day$ covariate is 0 for naïve mice, this covariate captures neural activity modes (bases) that are unique to trained mice. On each Test day, there were 600 trials, so trial number could range from 1 to 600. In addition, stimuli were presented in blocks of 50 trials, so the trial within block number could range from 1 to 50. The negative logarithm captures our intuition that adaptation and/or synaptic depression will generally have the effect of decreasing firing rates, though the unit factors allow for any given unit to be modulated in either direction. Significant interactions were: angle squared with a final rank of 1, E by angle with a rank of 1, E by angle squared with a rank of 1, second element held by day with a rank of 1, second element held by angle with a rank of 1, E by day with a rank of 1, trial number squared with a rank of 2, and trial number by trial within block with a rank of 1. We included all other possible interaction terms in the model fitting procedure, but all had a final rank of 0. We also included one triplet interaction term: E by second element held by day, which had a final rank of 0.

**Supplemental Table 2: Summary of Statistical Tests**

| Figure | Time Window | Data Mean | Data Standard Deviation | Approximate 95% Confidence Interval | Test Type | Test Statistic |
|---|---|---|---|---|---|---|
| 3b | Early | Day 1 FR· 0.453<br>2· 0.377<br>3· 0.384<br>4· 0.410 | 1· 0.434<br>2· 0.346<br>3· 0.305<br>4· 0.403 | 1· [0.31,0.60]<br>2· [0.25,0.50]<br>3· [0.28,0.49]<br>4· [0.27,0.55] | Two-sided permutation test | Difference in mean FR from Day 1 (naïve) |
| 3b | Late | 1 FR· 0.149<br>2· 0.023<br>3· 0.028<br>4· -0.027 | 1· 0.160<br>2· 0.139<br>3· 0.202<br>4· 0.281 | 1· [0.09,0.20]<br>2· [-0.03,0.07]<br>3· [-0.04,0.10]<br>4· [-0.12,0.07] | Two-sided permutation test | Difference in mean FR from Day 1 |
| 4d | Early | 1 FR Difference· -0.200<br>2· -0.040<br>3· -0.066<br>4· -0.104 | 1· 0.392<br>2· 0.276<br>3· 0.262<br>4· 0.198 | 1· [-0.33,-0.07]<br>2· [-0.14,0.06]<br>3· [-0.16,0.02]<br>4· [-0.17,-0.04] | Two-sided permutation test | Difference in mean FR Difference from Day 1 |
| 4d | Late | 1 FR Difference· -0.005<br>2· 0.153<br>3· 0.106<br>4· 0.155 | 1· 0.189<br>2· 0.201<br>3· 0.200<br>4· 0.223 | 1· [-0.07,0.06]<br>2· [0.08,0.23]<br>3· [0.04,0.17]<br>4· [0.08,0.23] | Two-sided permutation test | Difference in mean FR Difference from Day 1 |
| 5d | Early | 1 FR Difference· -0.169<br>2· -0.007<br>3· -0.084<br>4· -0.009 | 1· 0.254<br>2· 0.234<br>3· 0.346<br>4· 0.254 | 1· [-0.25,-0.84]<br>2· [-0.09,0.08]<br>3· [-0.20,0.03]<br>4· [-0.10,0.08] | Two-sided permutation test | Difference in mean FR Difference from Day 1 |
| 5d | Late | 1 FR Difference· 0.006<br>2· 0.095<br>3· 0.057<br>4· 0.182 | 1· 0.136<br>2· 0.195<br>3· 0.288<br>4· 0.183 | 1· [-0.04,0.05]<br>2· [0.02,0.17]<br>3· [-0.04,0.15]<br>4· [0.12,0.24] | Two-sided permutation test | Difference in mean FR Difference from Day 1 |
| 6b | Early ($A\tilde{B}$) | 1 FR· 0.252<br>2· 0.174<br>3· 0.211<br>4· 0.267 | 1· 0.294<br>2· 0.315<br>3· 0.249<br>4· 0.310 | 1· [0.15,0.35]<br>2· [0.06,0.29]<br>3· [0.13,0.30]<br>4· [0.16,0.37] | Two-sided permutation test | Difference in mean FR from Day 1 |
| 6b | Late ($A\tilde{B}$) | 1 FR· 0.200<br>2· 0.082<br>3· 0.052<br>4· -0.002 | 1· 0.264<br>2· 0.253<br>3· 0.245<br>4· 0.255 | 1· [0.11,0.29]<br>2· [-0.01,0.17]<br>3· [-0.03,0.14]<br>4· [-0.09,0.8] | Two-sided permutation test | Difference in mean FR from Day 1 |
| 6b | Early ($E\tilde{B}$) | 1 FR· 0.418<br>2· 0.411<br>3· 0.320<br>4· 0.289 | 1· 0.388<br>2· 0.483<br>3· 0.375<br>4· 0.342 | 1· [0.29,0.55]<br>2· [0.23,0.59]<br>3· [0.19,0.45]<br>4· [0.17,0.41] | Two-sided permutation test | Difference in mean FR from Day 1 |
| 6b | Late ($E\tilde{B}$) | 1 FR· 0.144<br>2· 0.045<br>3· 0.045<br>4· 0.039 | 1· 0.237<br>2· 0.245<br>3· 0.269<br>4· 0.307 | 1· [0.06,0.22]<br>2· [-0.04,0.13]<br>3· [-0.05,0.14]<br>4· [-0.07,0.14] | Two-sided permutation test | Difference in mean FR from Day 1 |

| | | | | | | |
|---|---|---|---|---|---|---|
| 7c – Real Data | n/a | 1 Soft Accuracy· 0.260<br>2· 0.245<br>3· 0.298<br>4· 0.326 | n/a | 1· [0.241,0.278]<br>2· [0.227,0.263]<br>3· [0.279,0.317]<br>4· [0.306,0.346] | Two-sided permutation test | Difference in soft accuracy from Day 1 |
| 7c – Simulated Data | n/a | 1 Soft Accuracy· 0.594<br>2· 0.591<br>3· 0.648<br>4· 0.644 | n/a | 1· [0.574,0.615]<br>2· [0.570,0.612]<br>3· [0.627,0.668]<br>4· [0.623,0.664] | Two-sided permutation test | Difference in soft accuracy from Day 1 |

n/a is not applicable
